# Streamlined freshwater bacterioplankton *Nanopelagicales* (acI) and “*Ca*. Fonsibacter” (LD12) thrive in functional cohorts

**DOI:** 10.1101/2020.03.18.997650

**Authors:** Rhiannon Mondav, Stefan Bertilsson, Moritz Buck, Silke Langenheder, Eva S. Lindström, Sarahi L Garcia

## Abstract

While fastidious microbes can be abundant and ubiquitous in their natural communities, many fail to grow axenically in laboratories due to auxotrophies or other dependencies. To overcome auxotrophies these microbes rely on their surrounding cohort. A cohort may consist of kin (ecotypes) or more distantly related organisms (community) with the cooperation being reciprocal or non-reciprocal, and expensive (Black Queen hypothesis) or costless (byproduct). These metabolic partnerships (whether at single species population or community level) enable dominance by and coexistence of these lineages in nature. Here we examine the relevance of these cooperation models to explain the abundance and ubiquity of the dominant fastidious bacterioplankton of a dimictic mesotrophic freshwater lake. Using both culture dependent (minimalist mixed cultures) and culture independent (SSU rRNA gene time series and environmental metagenomics) methods we independently identified the primary cohorts of *Actinobacterial* genera “*Ca*. Planktophila” (acI-A) and “*Ca*. Nanopelagicus” (acI-B), and the *Proteobacterial* genus “*Ca*. Fonsibacter” (LD12). While “*Ca*. Planktophila” and “*Ca*. Fonsibacter” had no correlation in their natural habitat, they have the potential to be complementary in laboratory settings. We also investigated the bi-functional catalase-peroxidase enzyme KatG (a common good which “*Ca*. Planktophila” is dependent upon) and its most likely providers in the lake. Further we found that while ecotype and community cooperation combined may explain “*Ca*. Planktophila” population abundance, the success of “*Ca*. Nanopelagicus” and “*Ca*. Fonsibacter” is better explained as a community byproduct. Ecotype differentiation of “*Ca*. Fonsibacter” as a means of escaping predation was supported but not for overcoming auxotrophies.

**IMPORTANCE:** This study examines evolutionary and ecological relationships of three of the most ubiquitous and abundant freshwater bacterial genera: “*Ca*. Planktophila” (acI-A), “*Ca*. Nanopelagicus” (acI-B), and “*Ca*. Fonsibacter” (LD12). Due to high abundance, these genera might have a significant influence on nutrient cycling in freshwaters worldwide and this study adds a layer of understanding to how seemingly competing clades of bacteria can co-exist by having different cooperation strategies. Our synthesis ties together network and ecological theory with empirical evidence and lays out a framework for how the functioning of populations within complex microbial communities can be studied.

## INTRODUCTION

The stable coexistence of species is described by a continuum of inter-specific symbiotic interactions from parasitism through neutralism to mutualism (1). The extended coexistence of microbial strains is described by intra-specific relationship models limited to niche theory (ecotype differentiation) (2) and predator control of dominant strains (3). Combining these ecological theories of co-existence with phylogenetic information can produce a simplified model with a new holistic perspective for understanding aquatic microbial communities. In aquatic microbial communities, especially for non-particle-associated bacterioplankton, physical structures and proximity as seen in soil, host-3 associated microbiome, and biofilm communities (4) are absent. As cooperation models have been developed from the study of such tightly interwoven assemblages their models may not be the most suitable. Absent also is the habitat isolation which limits dispersal as seen in e.g. soil ecosystems. In aquatic systems interactions between nutrients and biota can occur from micro up to macro-scale distances involving stochastic effectors from Brownian motion up to whole system circulation currents. Direct but distant interactions are therefore difficult to identify based purely on proximity as common goods and toxins in the environment can come from multiple sources and be transported long distances. Data from long-term monitoring of aquatic environments can be used to test the validity of ecological models.

Lake Erken has been extensively studied for several decades, and ample background information on nutrient cycling, planktonic communities and biogeochemical processes combined with well-established infrastructure makes this ecological observatory an excellent choice for the study of competing microbial assemblage processes (5–7). Several studies have investigated the microbial component of this lake ecosystem (8–13), revealing that the most abundant microbes in Lake Erken belonged to the LD12 (now described as the candidate genus “*Ca*. Fonsibacter” (14)) and acI (ACK-M1, hgcI, but now described as the candidate order *Nanopelagicales*) clades (11, 12, 15–20). “*Ca*. Fonsibacter” is a non-marine genus within the ubiquitous and abundant *Pelagibacteraceae* family which includes the marine *Pelagibacter* genus (SAR11). *Nanopelagicales* is a freshwater order with two described genera, “*Ca*. Planktophila” and “*Ca*. Nanopelagicus”. Both “*Ca*. Fonsibacter” and the *Nanopelagicales* have small cell size, reduced genomes (1.16 to 1.48 Mbp), multiple auxotrophies, a requirement for reduced sulfur compounds, have rhodopsins, and have been recalcitrant to maintenance in axenic laboratory culture (14, 21–23). “*Ca*. Fonsibacter” and “*Ca*. Nanopelagicus” genera both have additional amino-acid auxotrophies (23, 24). Recently a single strain of “*Ca*. Fonsibacter” and two strains of the *Nanopelagicales* genus “*Ca*. Planktophila” (acI-A and acI-A4) were successfully maintained in axenic cultures by customized liquid media in the case of “*Ca*. Fonsibacter” (14) and the addition of an active enzyme to filtered lake-water media for “*Ca*. Planktophila” (25). While both have been cultured neither have been deposited in culture collections and so retain *Candidatus* status. The difference in culture methods point towards a difference in lifestyle strategies and therefore evolution and ecology in these two abundant and ubiquitous freshwater bacterioplankton clades.

The ubiquity of the small-genome bacterial lineages, such as *Nanopelagicales* and “*Ca*. Fonsibacter” in freshwaters, is thought to be a consequence of their streamlined genomes and small cell size (26). On one hand, small cell size increases surface area to volume ratio, and improves diffusion, therefore decreasing time for foraging for non-motile organisms (4). On the other hand, reduced genomes have lost extraneous metabolic functions while having very low gene redundancy, reduced regulatory components, and high coding density (27). This combined streamlining and shrinkage reduces the energetic and material cost of genome, proteome, and cellular maintenance. Because their small genomes encode fewer metabolic functions than average, it has been proposed they require a large effective population (large *N_e_*) within which ecotypes diverge to provide the genomic flexibility usually seen in individual cells of bacteria with larger genomes. A large *N_e_* is also required for purifying selection to be efficient at reducing and maintaining small genome size. Such streamlined microbes were first noted in low nutrient aquatic environments and streamlining has since then been suggested as being of selective advantage under reduced nitrogen availability (28). *Nanopelagicales* and “*Ca*. Fonsibacter” clades are dominant and successful in non-marine environments such as Lake Erken, which far from being nutrient limited is classified as mesotrophic on a eutrophication trajectory. Further, as Lake Erken is ice and snow covered most winters (temperature and light limiting) and stratified in summer (thermo and oxy-clines), it is not a stable environment where organisms with reduced regulatory capacity are expected to thrive. It is unlikely that dominance in such productive lakes is made possible purely by streamlining, as the subsequent loss of genomic flexibility required to respond to a dynamic and variable habitat such as Lake Erken would counteract the benefits. Furthermore, large populations attract predators though both clades may escape some grazing predation due to their small size (29, 30). “*Ca*. Fonsibacter” may additionally have similar membrane properties to its sister marine lineage SAR11(31) to further elude grazing. All three genera encode cell membrane modification genes in hypervariable regions predicted to assist escape from viral predation (23, 32). A large *N_e_* where the divergence is focused in immunogenic or predator-escape mechanisms (defense specialist) will help a species survive population sweeps caused by predators, though it does not explain the intra nor inter-specific diversity or the seeming lack of competition between these three genera.

We combined the continuum of symbiotic interactions with phylogenetic information to produce a simplified model with a new holistic perspective for understanding aquatic microbial communities. (Fig. 1). Ecotype cooperation model describes how anabolic variability within a species population i.e. strain level variation, enables all the strains working together to produce nutrients required for cellular function and replication. It also allows for recombination and swapping of metabolic modules across strains. The community cooperation model also has a basis in anabolic complementarity but places the metabolic variation within a symbiotic group of different species whereby altruistic members provide expensive (Black Queen Hypothesis) or costless common goods (33, 34). The community detrital model is similar to community cooperation but focuses on uptake and catabolism of nutrients released by cellular death mediated by toxicity, starvation, or predation. Aquatic habitats in particular, due to transport and diffusion, are understood to have a unique niche for detritivores that may select for streamlined microbes (35).

**Fig. 1.**
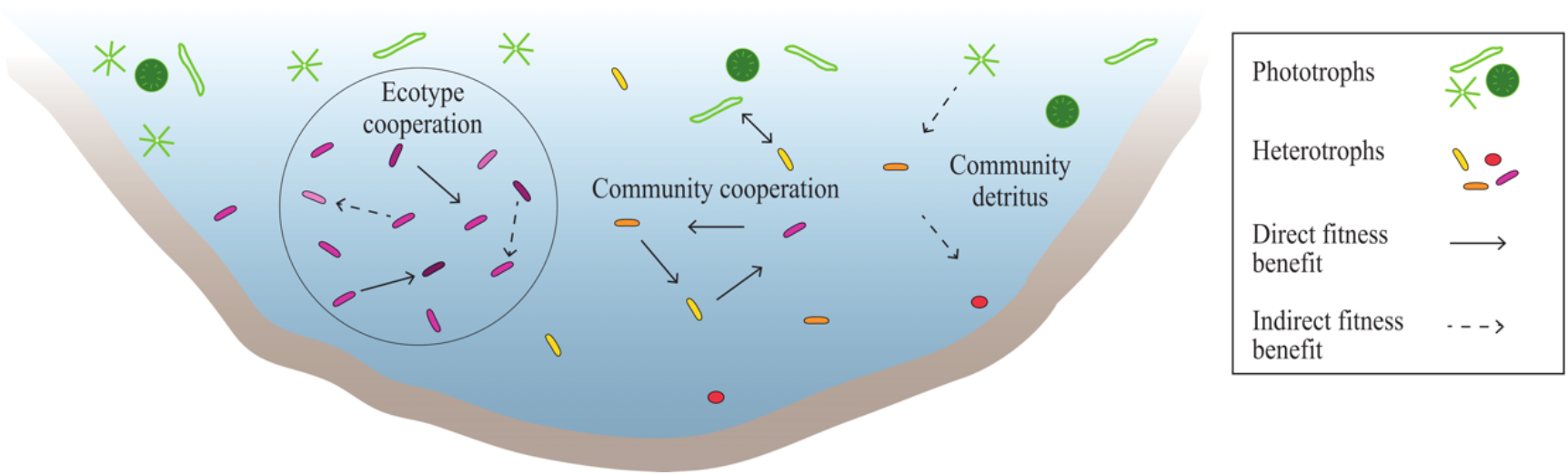
Cooperation models shown in a lake setting. Ecotype Cooperation Model describes anabolic variability as the main contributor to co-existence while allowing for recombination and swapping of metabolic modules across strains. Community Cooperation Model also encompasses anabolic complementarity but between rather than within species and includes common goods ranging from expensive to costless. The Community Detrital Model places complementarity both between and within species but describes catabolic variation.

Here we examine an 8-year time-series along with 26 minimalist mixed-cultures and selected lake metagenomes, to identify putative supporting microbes (cohort) and whether that support is best described by ecotype, community, and/or detrital models. We began with OTU network modelling (to define a candidate cohort that might provide for the target three genera dependencies), examined the phylogenetic relatedness of each cohort (to determine if a signal for ecotype or community existed), reviewed the metabolic diversity of the target group (to establish its dependencies and diversity), and investigated potential providers of the common-goods catalase-peroxidase (KatG) (a known dependency of some *Nanopelagicales*) within the metagenomes. Our proposed framework focusses on neutral to positive interactions as the laboratory experimental design does not permit detection of negative interactions even though timeseries can expose potential negative interactions.

## RESULTS

### Taxon abundance and environmental correlates

The most abundant clades detected in the SSU rRNA gene 8-year time-series were the *Actinobacterial* order *Nanopelagicales* at 29% average relative abundance (av.ra), the eukaryote phytoplankton genus *Stramenopiles* 12% av.r.a, the *Actinobacterial* clade ‘acIV’ (C111) 11% av.ra, the *β-proteobacterial* family *Comamonadaceae* 5% av.ra, the *α-Proteobacterial* candidate genus “*Ca*. Fonsibacter” 4% av.ra, and the *Verrucomicrobial* family *Cerasicoccaceae* 3% av.ra (Fig. S1, Table S1). There were few significant correlations or differences (Pearsons correlation or Kruskal-Wallis (KW) with post-hoc testing to identify which pairs were significantly different (KWmc)) between dominant clade abundance and a broad range of environmental predictors such as lake-cycle/season or physico-chemical conditions of the water (Fig. S2, Table S2). The three significant linear correlations detected had only moderate strength (r >= 0.4) with cloud or wedge-shaped distributions (Fig. S3).

### Microbial networks

The OTU network from both the time-series and mixed-cultures (Fig. S4) shows modularity and taxonomic assortativity (grouping of related organisms or under-dispersion) at all taxonomic levels with greater assortativity in the mixed-cultures (Table S3). Assortativity increased with taxonomic level peaking at phylum level in the timeseries, while assortativity was lowest at phylum level in the cultures. First neighbours (the primary cohort) of OTUs in networks provide evidence (but not proof) for preferential association between phylotypes, or when negatively correlated may indicate mutual exclusion. The primary cohorts of “*Ca*. Planktophila, “*Ca*. Nanopelagicus”, and “*Ca*. Fonsibacter” (Fig. 2) are overlapping and include phylotypes from each other’s cohorts. Primary cohorts from the time-series and mixed-cultures are similar, though the correlation type and strength vary. There was a moderate correlation between relative abundance of OTUs in the timeseries (Fig. 2a) and relative abundance in cultures (Pearsons r = 0.58, padj = 0, Table S4) but no other correlations between relative abundance or prevalence (percentage of samples an OTU was detected in) were observed.

**Fig. 2.**
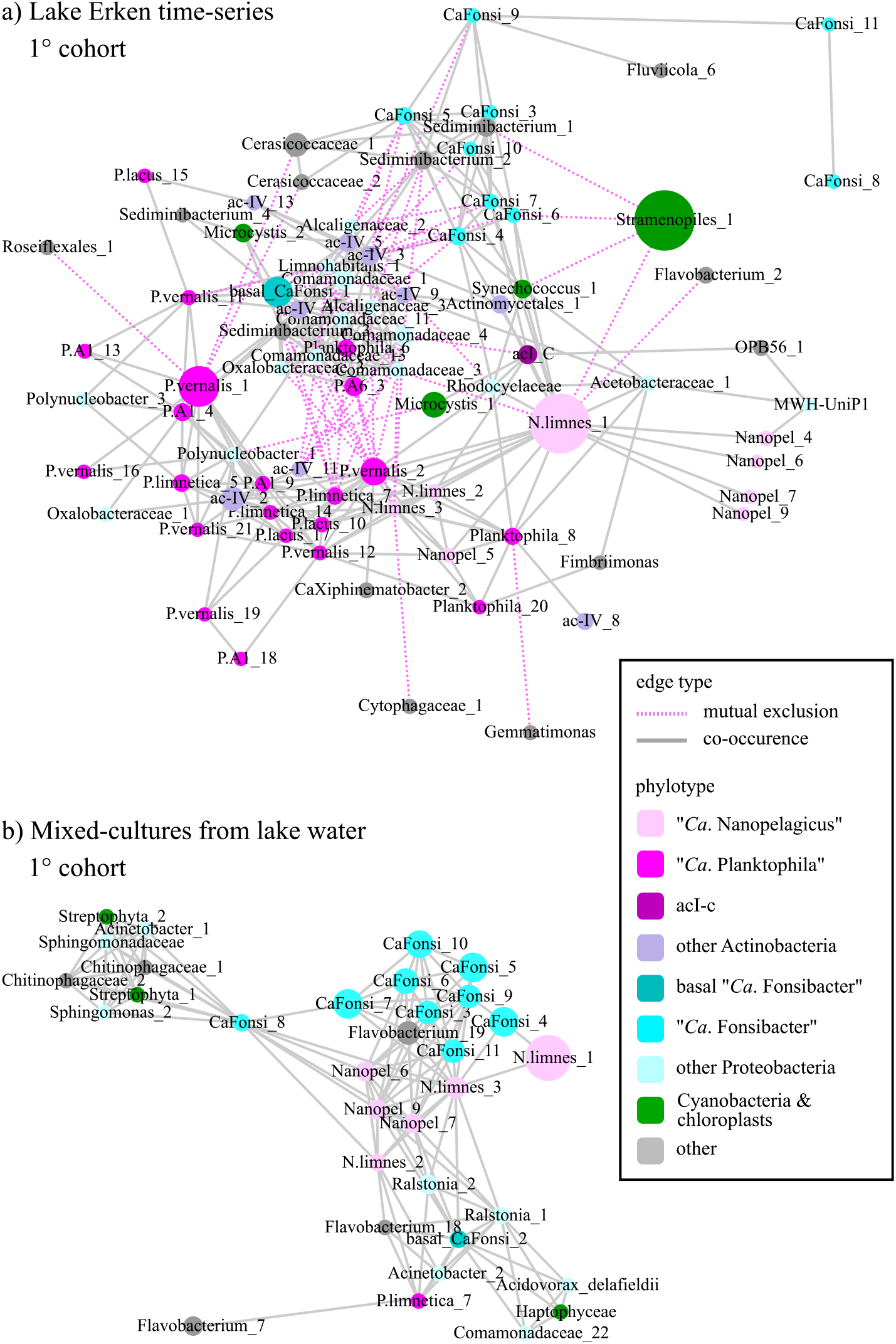
OTU networks of the primary cohorts from amplicons of a) time-series aand b) mixed-cultures. Nodes sized by average relative abundance in time-series and by percentage of cultures detected in for mixed-cultures. Pink dotted lines indicate mutual exclusion of OTUs while grey solid lines are co-occurrence.

### Actinobacterial order Nanopelagicales (acI)

*Nanopelagicales* in Erken had higher relative abundances in the hypolimnion, (KW p<0.001, KWmc p<0.001) and were negatively correlated with % O_2_ (Pearsons r = - 0.4, p<0.001). *Nanopelagicales* were the most abundant clade in Lake Erken and were represented by 31 OTUs in the lake time-series network with nine “*Ca*. Nanopelagicus”, 21 “*Ca*. Planktophila”, and one acl-c (Fig. 2a, Fig. S4). A subset of six “*Ca*. Nanopelagicus” and one “*Ca*. Planktophila” phylotype were detected in the mixed-culture based network (Fig. 2b, Fig. S4).

The time-series primary (1°) cohort (i.e. the cohort immediately surrounding) of “*Ca*. Planktophila” included: other *Actinobacteria, Armatimonadetes, Chloroflexi, Gemmatimonadetes, α-* and *β-Proteobacteria*, and *Verrucomicrobia*, while in the mixed-cultures only *Actinobacteria, Bacteroidetes*, and *α*- and *β-Proteobacteria* coexisted. A pairwise comparison of the phylogenetic distance and correlation (averaged edge value from the correlation network) was conducted to see if there was a relationship between OTU genetic relatedness and co-occurrence; to distinguish between ecotype and directed reciprocation models. Within the “*Ca*. Planktophila” 1° cohort, there was no support for a correlation between phylogenetic distance and cooccurrence (Fig. 3) and “*Ca*. Planktophila” was the only genera tested that had negative intra-genus correlations. “*Ca*. Planktophila” had more intra-phyla (i.e. non-*Nanopegicales Actinobacteria*) correlations than the other genera tested. Unlike the other genera, “*Ca*. Planktophila” had no correlations (positive or negative) with phytoplankton in its 1° network (Fig. 3). Additionally, “*Ca*. Planktophila” were the dominant Actinobacterial genus in the time-series with “*Ca*. P. vernalis” being most abundant (Fig. 2a). Conversely, in the mixed-cultures, only one *Planktophila* phylotype was detected, “*Ca*. P. limnetica” (Fig. 2b).

**Fig. 3.**
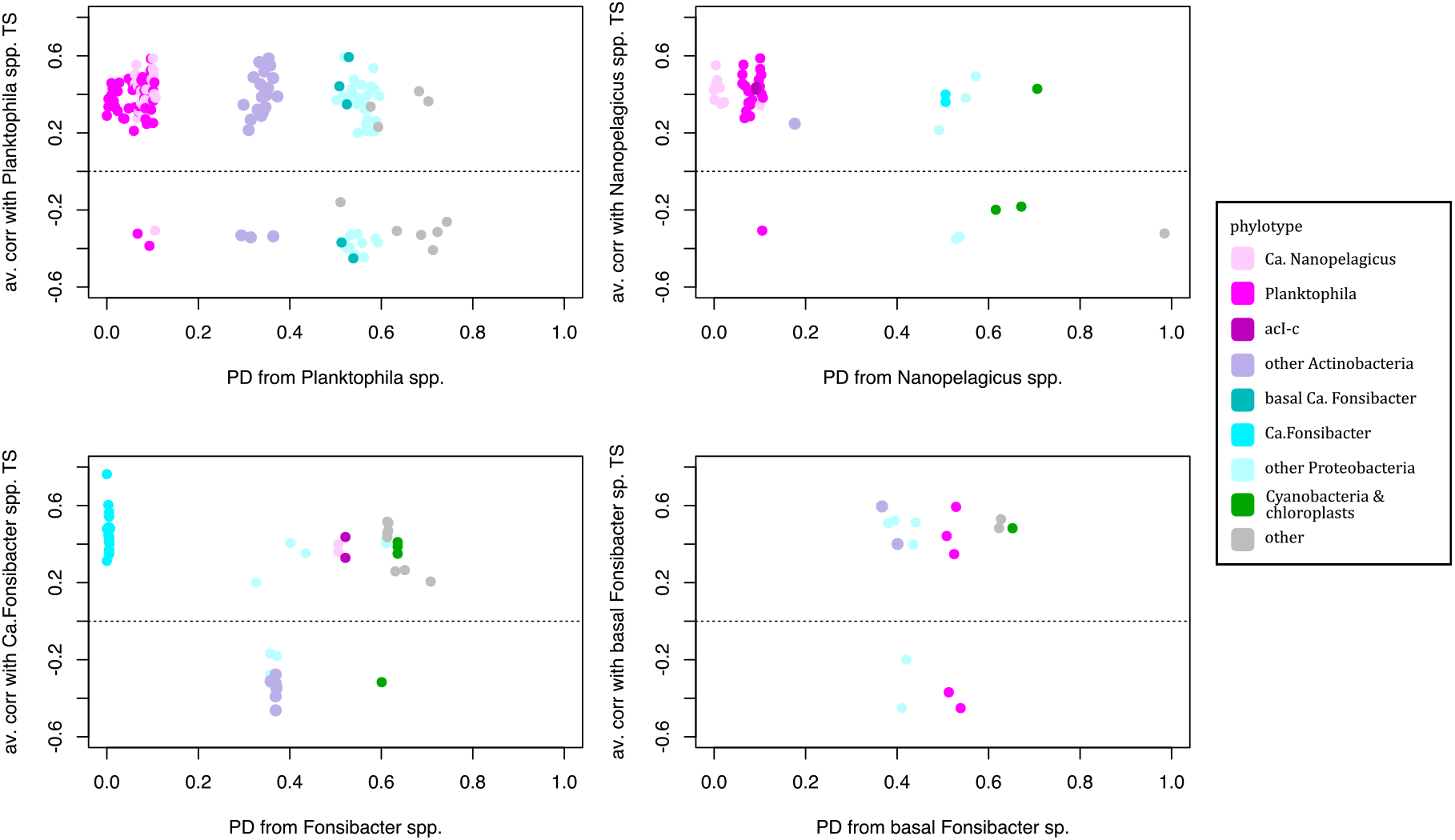
The relationship between OTU phylogenetic distance and correlation of the primary cohort phylotypes to selected genera: “*Ca*. Planktophila” upper-left, “*Ca*. Nanopelagicus” upper-right, “*Ca*. Fonsibacter” lower-left, and basal “*Ca*. Fonsibacter “lower-right.

“*Ca*. Planktophila spp.” were successfully maintained in axenic culture via the addition of bovine catalase and therefore the presence, absence, and functionality of the relevant *Nanopelagicales* catalase-peroxidase (KatG) is debated. We investigated the presence of the *katG* gene, its gene neighbourhood, and the homology and topology of its translated product, KatG, to establish if there is evidence to support its activity level or effect on host abundance. “*Ca*. P. vernalis” (acI-A7) KatG had highest homology to the high functioning *Synechoccocus* version (Fig. S5) and the gene is found near a tRNA (Fig. S6). “*Ca*. p. limnetica “s KatG had highest identity with the assayed low activity KatG from “*Ca*. P. rubra” (IMCC 25003) and was located within a putative DNA repair region (Fig. S5 & S6). When modelled these KatGs were most similar to *Burkholderiapseudomaleiis* with a predicted *β*-sheet around one of the active site residues instead of an *α*-helix, and are 10 to 20 amino-acid residues longer than known high functioning versions i.e. *Burkholderia* type A (Fig. S5). All known active site residues involved in heme interaction or enzyme switching were however present. “*Ca*. Planktophila limnetica“, “*Ca*. P. vernalis”, “*Ca*. P. lacus”, and “*Ca*. N. limnes” all have the repair cassette without *katG*. Metagenomic *katG* sequences were recovered matching “*Ca*. P. dulcis”, “*Ca*. P. limnetica”, “*Ca*. P. versatilis”, and “*Ca*. P. vernalis”.

“*Ca*. Nanopelagicus” 1° cohort in the time-series included other *Actinobacteria, Bacteroidetes, α*- and *β-Proteobacteria*, and the phytoplanktonic *Synechococcus* and *Stramenopiles* (identified by chloroplast sequences), while in the mixed-cultures there was a subset of these phyla with the addition of one *g-proteobacterial* phylotype (Fig 2). To calculate whether the presence of an OTU in the mixed-cultures could be attributed to the probability of it being in the inoculum, the total and average relative abundances, and the number of samples an OTU was detected in (prevalence) was compared across the two datasets with Pearsons correlation. There was no correlation between “*Ca*. Nanopelagicus” OTU relative abundance or prevalence in the lake time-series as compared to the mixed-cultures (Table S4). “*Ca*. Nanopelagicus” was overall more prevalent in mixed cultures than in the lake time-series. “*Ca*. Nanopelagicus” had strong correlation with “Ca. Fonsibacter” in both datasets. There were between 3 and 19 other phylotypes (non-*Nanopelgicales*, non-Fonsibacter) detected in cultures with “*Ca*. Nanopelagicus” (Table S5).

The most abundant “*Ca*. Nanopelagicus” phylotypes in both datasets were assigned to the *katG*^−^ “*Ca*. Nanopelagicus limnes”. Several metagenome-recovered putative “*Ca*. Nanopelagicus” *katG* genes were identified in IMG databases and while these KatGs had greatest homology with *Burkholderia* fold type B, they were not located within the repair-cassette and there was no synteny between them. No “*Ca*. Nanopelagicus” katG sequences were recovered from our metagenomes.

### α-Proteobacterial genus “Ca. Fonsibacter” (LD12)

“*Ca*. Fonsibacter spp.” had higher relative abundance from winter to spring and moderate negative correlation with water temperature (r = −0.4, p<0.001, Table S2). “*Ca*. Fonsibacter” was represented by 10 OTUs in each of the time-series and mixed-culture networks, with nine OTUs common to both and the alternating OTUs being the least abundant and least prevalent in both environments. The “*Ca*. Fonsibacter” 1° cohort from the time-series includes other “*Ca*. Fonsibacter”, *Bacteroidetes, α*- and *β-Proteobacteria, Actinobacteria* including “*Ca*. Nanopelagicus” and acI-c, and phytoplankton *Synechococcus* and *Stramenopiles*. The “*Ca*. Fonsibacter” 1° cohort from mixed-culture experiments also included other “*Ca*. Fonsibacter” phylotypes, “*Ca*. Nanopelagicus”, *Bacteroidetes*, other *α*- and *β-Proteobacteria* and macro-algal eukaryotes of the *Streptophyta* order. There was a strong correlation detected between “*Ca*. Fonsibacter” OTU prevalence in the lake time-series compared to the prevalence of the respective OTU in the mixed-cultures (corr = 0.82, n=11, p adj = 0, Table S4). Based on OTU identity, phylogenetic branch placement, and network connections, two “*Ca*. Fonsibacter” phylotypes were designated as basal and analysed separately for 1° cohort. Unlike the core

“*Ca*. Fonsibacter” phylotypes, these basal phylotypes had no negative correlations with phytoplankton and no correlations with other “*Ca*. Fonsibacter” phylotypes (Fig. 3).

### Other primary cohort members

Both the *Actinobacterial* ‘acIV’ (C111) and *Comamonadaceae* families were more abundant in winter when the lake was covered by ice (Fig. S2, KW p<0.001, KWmc p<0.001). While ‘acIV’ was abundant and ubiquitous in the time-series (Fig. S1, Table S1), and also prominent in the “*Ca*. Planktophila” time-series 1° cohort (Fig. 2a), no phylotypes were detected in the mixed-cultures. *Comamonadaceae* was abundant but patchy in the time-series and mixed cultures.

## DISCUSSION

### Taxon relative abundance and environmental correlates

If microbial abundance is directly linked to nutrient availability or other physico-chemical parameters, then a strong correlation between geo-chemical parameters and relative abundance is expected. The lack of strong correlation between OTUs or clades and environmental parameters in Lake Erken indicates that during the 8 years of sample collection, microbial community composition was not directly and strongly controlled, but only influenced, by the measured abiotic environmental variables. While short-term studies have shown significant strong trends and correlations (12), such correlations tend to be eclipsed in studies spanning multiple years by shifts operating at longer frequencies and biotic interactions (36–39). Overall, the strength and number (identified via consensus network) of correlations between taxon relative abundance and the weakness of environmental predictors for most of the taxa suggest that biological aspects (including predation, which we did not study) are the greater determinants of microbial population dynamics, composition and persistence in Lake Erken. It stands therefore that the dominance of the three streamlined genera in the lake is attributable more to their interactions with their community than selection by environment.

### Microbial networks

Assortativity is a predicted product of ecotype or niche differentiation at species level and environmental filtering at higher taxonomic levels (40). Ecotype divergence of strains reduces competition between different members of a population. The detection of highest assortativity in the time-series networks at phylum level supports environmental selection as important in the lake habitat but not at taxonomic levels lower than phyla. It is unlikely that environmental filtering is the main driver behind the success of these three genera in Lake Erken. High assortativity in the mixed cultures at all levels except phylum, supports that niche differentiation contributed to co-occurrence in the cultures. Neither result however fully supports, nor refutes, ecotype theory as the explanation of coexistence or dominance by “*Ca*. Planktophila”, “*Ca*. Nanopelagicus”, or “*Ca*. Fonsibacter” genera.

The strong correlation between “*Ca*. Fonsibacter” (and also *Comamonadaceae*) prevalence in the time-series and mixed-cultures supports that its occurrence in the cultures may be attributed to being present in the inocula and that the filtered lake water media was able to meet an important proportion of its metabolic needs. Conversely, the lack of correlation in either relative abundance or prevalence, for the *Nanopelagicales* genera (and also the ‘acIV’ family) in the time-series compared to mixed-cultures means that there is not support for their detection being merely due to high-abundance in the inocula. We infer that the co-presence of other phylotypes were bigger contributors to their growth. It should be noted that negative correlations are not detectable in the mixed-cultures as the absence of a phylotype could be stochastic rather than a result of competitive exclusion. Moreover, competition that prevents growth in a culture cannot be identified.

### Actinobacterial order Nanopelagicales ‘acI’

*Nanopelagicales* in Erken were likely more competitive in colder, darker, and oxygen depleted waters. While *Nanopelagicales* has been associated with oxygen saturated waters in some locations (41) this lineage has been associated with colder deeper water at others (30), though the variation in association could be due to the considerable genetic diversity encompassed in this bacterial order (42). It is also probable that solar-driven formation of oxygen radicals in surface waters inhibit the growth of *katG*^-^ species of *Nanopelagicales*.

Of the seven *Nanopelagicales* phylotypes in the mixed-cultures, six were members of the candidate genus “*Ca*. Nanopelagicus” suggesting that this genuss dependencies were more easily met compared to the “*Ca*. Planktophila spp.” found in Lake Erken. However, genomic evidence so far has shown the opposite as “Ca. Nanopelagicus spp.” have more numerous auxotrophies than “*Ca*. Planktophila spp.” (23). It is therefore possible that the difference in culturability seen in this experiment was due to reduced costs via gene loss in combination with suitable metabolic partners being in the inoculum. In particular the lower maintenance costs due to the loss of *katG* (25, 33) could increase fitness of the “*Ca*. Nanopelagicus”. The successful maintenance of “*Ca*. Planktophila” in culture pinpointed the low functionality of some *Nanopelagicales* catalase-peroxidases as the culprit for the difficulties in growing these bacteria. Peroxide is proposed to be an up-regulator of a DNA repair (43) and in the “*Ca*. Planktophila” with the putative low functioning KatG type, the gene is embedded in a region of DNA hosting other genes for repair of damage done to DNA under oxidative stress. *P. limnetica* (acI-A3) genomes have this repair cassette but their *katG* is located elsewhere. The most parsimonious explanation for *Nanopelagicales katG* presence/absence/homology is that the now low functioning version located in a repair cassette is in the process of being lost and that the “*Ca*. Planktophila spp.” with a high functioning KatG obtained this version via a HGT event. While *P. vernalis* in Lake Erken is likely *katG*^+^, it is likely that *P. limnetica* and *N. limnes* from Lake Erken rely on “*Ca*. Fonsibacter” in combination with either *Flavobacterium* or *Comamonadaceae* to detoxify peroxide. This would fit within the community cooperation model. While “*Ca*. Nanopelagicus” has more numerous auxotrophies compared to “*Ca*. Planktophila” (23) the higher culturability of “*Ca*. Nanopelagicus” supports that it may be the cost to habouring *katG*, whether high or low functioning, and as *katG* is larger than average (44), and as long as a functional aquaporin exists (45), and cells don’t grow too fast, then it would be an unnecessary cost, as cells can rely on other community members for oxidative stress reduction. Further the reduction in amino-acid synthesis pathways and therefore greater reliance on exogenous sources of amino-acids in “*Ca*. Nanopelagicus” appears to have paid off if their greater cultivability in these minimalist cultures is any indicator. This fits within the community detritus model.

### α-Proteobacterial genus “Ca. Fonsibacter” (LD12)

Earlier short-term studies found that “*Ca*. Fonsibacter” is highly abundant in Lake Erken in the late summer-early autumn immediately after an algal bloom i.e. the ’clear water’ phase of the lake’s cycle (12). However, we found support for higher long-term relative abundances in winter and spring and an associated negative correlation with water temperature. These differences in findings could be due to changes in abundance between years of different ecotypes (with different temperature and nutrient preferences) that were not detected via 16S surveys. The ecotype differentiation within “*Ca*. Fonsibacter” is centered around the HVR1 which is thought to encode genes for defense, i.e. predator escape via alteration of cellular membranes (3, 46). There appears to be limited intra-population variability in anabolic or catabolic traits (14, 24, 47) and as such, ecotype differentiation for overcoming auxotrophies is not supported. While no direct evidence exists for “*Ca*. Fonsibacter”, its similarity to the marine sister clade Pelagibacter suggests that it has overcome auxotrophic limitations by scavenging metabolites and other compounds produced by phototrophs (48). In Erken this would primarily be dying and senescent *Stramenopiles* as reflected in the apparent negative correlation. Evidence points to “*Ca*. Fonsibacter” relying on detritus to meet its dependencies and ecotype differentiation for defense.

The combination of network and other correlation analyses with review of published genomes supports that the most of *Nanopelagicales* have anabolic ecotype cooperation with reliance on a detoxifying community cohort as main structuring factors. A few “*Ca*. Planktophila” are putatively KatG+ but as they were not detected in Lake Erken it was not possible to determine the most appropriate cooperation models in this study. “*Ca*. Fonsibacter” may escape predation via ecotype differentiation, while broadly relying on community detritus for anabolic deficits.

## MATERIALS AND METHODS

### Time series water sampling and environmental parameters

Time series water samples were obtained from dimictic Lake Erken (59.1510N, 18.1360E) central Sweden at monthly to weekly intervals spanning a total period from February 2007 to November 2015 (Table S6). Samples for genomic and chemical analysis were taken from the 20 m water column at one meter intervals from the deepest point of the lake (Fig. S7). Samples for the total water column were pooled during periods of mixing. Samples of the upper oxygenated water column (epilimnion), middle (metalimnion), and lower oxygen depleted (hypolimnion) were pooled into separate composite samples during summer stratification. Annual dates of mixing, stratification, ice-over, and ice-thaw varied, as did depth of stratigraphic layers (Table S6). Temperature and oxygen concentrations were analyzed every meter with a portable Oxi 340i oxygen meter equipped with a Cellox 325-20WTW probe and were used to determine stratification. Chemical analysis of pooled water samples was carried out by the Lake Erken Field Station ISO certified laboratory and included pH, conductivity, nitrate, nitrite, alkalinity, turbidity, suspended matter, phosphate, ammonium, total and particulate phosphorus and nitrogen, chlorophyll-a, suspended particulate organic matter, silicate, water-colour and absorbance at 420nm measurements. Water samples for genomic analysis were collected by gentle vacuum filtration onto 0.2 μm membrane filters (Supor-200 Membrane Disc Filters, 47mm; Pall Corporation, East Hills, NY, USA). Filters were individually stored in Eppendorfs and transported back to the University Uppsala laboratory where they were stored at −80 °C until further processing.

### DNA extraction and SSU rRNA gene amplicon preparation

DNA was extracted from filters using MoBio Ultra-clean soil DNA extraction kits (MoBio, Carlsbad) as per manufacturers’ instructions. The V3-V4 region of the 16S rRNA gene was amplified using the S-D-Bact-0341-b-S-17 (Bakt_341F: CCTACGGGNGGCWGCAG) forward primer and S-D-Bact-0785-b-A-21 (805RN: GACTACNVGGGTATCTAATCC) reverse primer (49). Template DNA was amplified in duplicate 20 μl reactions containing 1.0 U Q5 high fidelity DNA polymerase (NEB, UK), 0.25 μM primers, 200 μM dNTP mix, and 0.4 μg BSA. The thermocycler program was an initial denaturation step at 95 °C for 30 s, followed by 20 cycles of dissociation at 95 °C for 10 s, annealing at 53 °C for 30 s extension at 72 °C for 30 s, with a final extension of 2 min at 72 °C. Amplicons were pooled and purified with Agencourt AMPure XP purification system (Beckman Coulter, Danvers, MA, USA). Purified amplicons (2 μl) were amplified in second step 20 μl reactions to introduce sample MIDS to each end of the amplicons. Reactions contained 1.0 U Q5 high fidelity DNA polymerase (NEB, UK), 0.25 μM primers, 200 μM dNTP mix, and 0.4 μg BSA. Thermocycler program was an initial denaturation step at 95 °C for 30 s, followed by 15 cycles of dissociation at 95 °C for 10 s, annealing at 66 °C for 30 s, extension at 72 °C for 30 s, with a final extension of 2 min at 72 °C. Amplicons were again purified using Agencourt AMPure XP purification system (Beckman Coulter, Danvers, MA, USA) then quantified using PicoGreen (Invitrogen) before pooling at equi-molar amounts for each run. Amplicons were sequenced at SciLifeLab SNP/SEQ service on the MiSeq Illumina platform with 300bp paired-end libraries. MiSeq data was processed, including de-multiplexing, by the sequencing centre using Illumina pipelines where all reads with more than 8% mismatch to adapter-MID sequences were discarded.

### Water sampling, culture conditions, and DNA amplification for mixed-cultures

Water as both a source of growth-media and culturable-cells was collected from the epilimnion of Lake Erken on March 8^th^ 2016. Water for growth-media was filter-sterilised twice through Sterivex filter cartridges 0.22 μm (Millipore). Ten minutes exposure to UV light further disinfected and disrupted cellular and viral nucleic acids. Ten microliters of untreated lake water (10^6^ cells/ml) were used as a source of cells for inoculation of mixed-cultures. The untreated lake water was diluted in 1 L of the triple sterilized lake water (approximately 10 cells/ml). Ninety-eight 1 ml cultures were placed in sterile 50 ml falcon tubes to allow oxygenation and incubated for two months under 12-hour light-dark cycles at 11 °C to replicate lake conditions at time of collection. The V3-V4 region of the 16S rRNA gene was amplified using the S-D-Bact-0341-b-S-17 (Bakt_341F: CCTACGGGNGGCWGCAG) forward primer and S-D-Bact-0785-b-A-21 (805RN: GACTACNVGGGTATCTAATCC) reverse primer (49) One microliter of the mixed-culture after one freeze-thaw cycle was used as template and Q5^®^ High-Fidelity DNA Polymerase. The PCR conditions consisted of an initial step at 98°C for 10 minutes.

Thereafter, 35 cycles of denaturation at 98°C for 10 s, annealing for 30 s and extension at 72°C for 30 s. Annealing temperature was 48°C which has been tested to get unbiased products of non-mismatch and 3-mismatch isolates. Final extension was performed at 72°C for 2 minutes. PCR products were purified with magnetic beads (Angecourt AMPure). A second PCR was conducted for attaching standard illumina handles and index primers as described above. Primers and MIDS detailed in Table S7. Prior to sequencing a portion of amplification product was run on gel to check for amplification and used as proxy for growth of culture. Only data from the 60 cultures with a visible band after the first PCR were included in analyses.

### Amplicon bioinformatics

Demultiplexed MiSeq time-series and mixed-culture data, and 454 Ti time-series data obtained from Lake Erken 2008 (11), were pre-processed separately and joined once chimeras had been removed. Pre-processing of MiSeq data started with SeqPrep v1.2 (https://github.com/jstjohn/SeqPrep.git) with default settings to join paired end reads while discarding reads shorter than 30b and all un-paired reads. 454 and MiSeq reads with ambiguous bases, low quality average, and low quality sections, were discarded and trimmed with Qiime v1.9 (50) split_libraries and multiple split libraries fastq respectively. Singletons and sequences identified as chimeric were removed with Qiimes’ identify_chimeric_seqs, filter_fasta using usearch v6.1 (51), with default settings and option parameter ‘intersection’ to select out sequences identified by both de-novo and reference-based methods. Both GreenGenes (v105 13-8) and SILVA (v128) reference datasets were used (52, 53). Sequences from the two Miseq runs and the 454 data from 2008 were then combined. Reads were clustered at 97% identity into OTUs then assigned taxonomy based on SILVA v128 using QIIMES closed pick OTU method (uses only template recognition and does not allow for de-novo sequences) to allow for differences in DNA processing and sequencing across the three datasets. Sequences from the negative control were removed from the entire dataset. Samples with potential human-microbiome contamination (Table S8) were identified and discarded from the dataset due to potential effect on community metabolism. Sequences identified as eukaryotic (including fungal, mitochondrial and chloroplast origin) or archaeal were deliberately retained despite the primers not specifically being selected to amplify such sequences. as they were considered critical in the mixed-culture analyses from a community metabolism perspective. Samples were normalized using Qiimes single rarefraction to even depth of 1500 reads per sample for the time-series and 500 reads for the mixed-culture samples. These normalised OTU tables were then used for all downstream analyses.

### Statistical analyses of time-series environmental correlates and selected taxa

Statistical analyses were done using R v3.3.1 (54) in the RStudio IDE v0.99.903 (55) and graphs processed for publication in Inkscape v0.91. Environmental parameters were removed if more than 40% of the values were missing then remaining twenty-one tested for co-linearity with the usdm v1.1.18 package with vifstep followed by vifcor functions and removed if VIF >10 or correlations were over 0.7 sequentially starting with those with highest number of missing values. The remaining 14 environmental predictors were tested for statistical differences in connection to lake-cycle with Kruskal-Wallis (kruskal.test of the base stats package) and graphed using beanplot v1.2 (56) and scales v0.4.1. Statistical significance of differences between individual paired cycle points were determined post-hoc with the Kruskal-Wallis multiple comparison (KWmc) using thekruskal.mc test of the pgirmess package v1.6.4 (57). Environmental parameter correlates with selected taxa relative abundances were screened using Pearsons correlation with an alpha level of 0.001 and p-values estimated and corrected for multiple comparisons with Benjamini, Hochberg, and Yekutieli (BH) false discovery rate using the psyche package v1.6.12 corr.test function and Pearsons moment r reported. Parameters with weak to moderate correlation (r >|0.3|) were plotted in R to observe monotonicity.Network analyses

The normalized time-series OTU table was further processed for network and correlation analyses to remove rare OTUs that, while potentially important within the community, are known to inhibit identification of ecological relationships between OTUs and also between OTUs and their environment. Recommendations include OTU tables with 50% or less sparcity and neff (inverse Simpsons) above an average of 10 and the use of multiple methods (58). A range of filtering parameters based on frequency of OTU detection, from 2 samples up to 46 (20% of samples) and total relative abundance ranging from 1 up to 150 (0.1%), was used on the OTU table to remove type I errors in detection of co-occurrence networks (58). The best filter combination was found with OTUs at a total relative abundance above 0.1% and occurring in at least 29 samples. All OTU tables were analysed in R (54) using the vegan v2.4-1 package (59) diversity function to find the neff and the base stats v3.3.1 package to find min, max and mean of the neff. The reduced OTU table was analysed with SparCC (60), Pearsons correlation coefficient, and SPIEC-EASI (61). SparCC analyses were carried out in Rstudio (55) using both the rsparcc and spiec.easi packages with rsparcc parameters: max.iter=100, th=0.1, exiter=10 and pseudo p-values generated with spiec.easis sparccboot (100 bootstraps) and pval.sparccboot scripts (two-sided test). Pearson correlations were performed using the corr.test of psyche package using the optional p-values adjustment for multiple testing of Benjamini, Hochberg and Yekutieli false discovery rate. SPIEC-EASI correlations were analysed using the spiec.easi package with parameters: method=“mb”, sel.criterion=“stars”, lambda.min.ratio=1e-2, nlambda=20, and 50 repetitions. Singletons were removed from the normalized mixed-culture OTU table prior to network analysis. The reduced OTU table was analysed with Pearsons correlation, SpiecEasi (61) and on the binary version of the table, Dice-Sørensen index (62, 63).

Correlations between OTUs in the time-series and mixed cultures were retained if the correlation co-efficient was greater than 0.3 absolute for Pearsons, SparCC, or Dice-Sørensen, or if detected at all by SPIEC-EASI. Time-series and culture networks consisting of OTU connections detected by at least two correlation methods were visualized in Cytoscape. All further statistical analyses of the networks were performed in Cytoscape except for assortativity analyses which were done in R using the igraph v1.1.2 package nominal assortativity function. Sub-networks of the identified dominant genera (“*Ca*. Planktophila”, “*Ca*. Nanopelagicus”, and “*Ca*. Fonsibacter”) and their primary cohorts were extracted from the two initial networks and visualized separately. Layout for the networks were “edge-weighted spring-embedded” with edge weights calculated from the average of the correlations. Some nodes (OTUs) were moved to the side of overlapping nodes for clarity or away from an edge if it appeared to pass through an unconnected node. Correlations between *Nanopelagicales* OTU and “*Ca*. Fonsibacter” OTU detection in the time-series versus the mixed cultures were investigated by Pearson correlation of their relative abundance (log av.ra), prevalence (percent samples or cultures detected in) and relative abundance versus prevalence to determine if detection in culture was correlated to occurrence in the lake.

### Metagenomic sequencing

Samples for metagenomic sequencing were selected by highest relative abundance of clades of interest according to amplicon data, greatest available DNA mass, and greatest variation in environmental parameters. This was done to examine samples with the greatest potential divergence of environmental conditions under which microbes of interest were most abundant. These included samples taken during lake circulation and summer stratification which separately including hypo, meta and epi-limnionic samples. Two samples per category were selected from different years. Lower biomass during ice cover meant there was insufficient DNA for metagenomic sequencing. DNA library for metagenomic sequencing was prepped using the Illumina TruSeq Nano, 550bp NeoPrep and sequenced with Illumina HiSeq 2500 High Output V4 PE 2×125bp at SciLifeLab Stockholm.

### Metagenome bioinformatics

HiSeq Illumina paired end reads were pre-processed first using SeqPrep v1.2 (64) to filter out low quality reads using default settings resulting in removal of all reads less than 30b in length, un-paired reads and initial clipping of adapter sequences. Next sequences were cleaned using Trimmomatic v0.36 (65) with removal of any remaining Illumina adapters allowing for 2 mismatches; followed by 4b sliding window trim with minimum average quality of 15; then trimming of read ends with ‘maxinfo’ value less than 1. The eight metagenomes were individually assembled with megahit (66) using iterative kmers from 21 to 121 at 10b intervals, minimum coverage of 2, and minimum length of 200. Concatenated assemblies were cleaned of Illumina artifact low complexity (long homopolymer) contigs with prinseq-lite v0.20.4 (67) removing contigs consisting of greater than 80% one nucleotide.

### Investigation of katG gene and its product: catalase-peroxidase I

Known impediments to axenic growth of freshwater isolates are multiple auxotrophies, and slow growth and replication rates. The successful axenic cultivation of “*Ca*. Planktophila” strains by addition of high functioning catalase to reduce oxidative stress, in addition to catering for predicted auxotrophies, showed that there can be other considerations. To investigate how both “*Ca*. Planktophila” and “*Ca*. Nanopelagicus” phylotypes grow in Lake Erken and why “*Ca*. Nanopelagicus” grew more frequently in the minimalist mixed cultures, the katG gene was investigated. First katG amino-acid sequences for species identified as belonging to the Nanopelagicales and its primary cohort were retrieved from IMG and GenBank (68, 69). Second sequences for whom the gene product has been assayed and/or the 3D crystal structure is known were identified via Phyre2 and then collected from RCSB PDB (rscb.org (70, 71)). Next these sequences were used as BLAST (72) queries to retrieve closely related (> 80% aa identity) sequences from Lake Erken metagenomes. The collected amino-acid sequences were aligned using MAFFT e-insi and “leavegappy” options (73). Sequences with greater than 99% identity were identified using CD-Hit and removed (74) while sequences from uniRef useful for constructing a guide tree were then also added (75, 76). The alignment was evaluated for best amino-acid substitution model using IQ-tree v.1.6.12 modelfinder (77). This alignment was then analysed and used to construct a tree in RAXML v.0.9.0 (78) with WAG+IG+R4 model and 1000 bootstraps and catalase-peroxidase from *Oryza rufipogon* set as outgroup. To examine functionality secondary structure was predicted and aligned to known models then tertiary structure predicted in Phyre^2^ (71). Further the location of katG genes in Nanopelagicales genomes and MAGs was examined in IMG using the gene-neighbourhood function. Images were downloaded with screen shot and processed in Inkscape.

### Data availability

Time-series MiSeq sequences were deposited in ENA (https://www.ebi.ac.uk/ena/browser/view/PRJEB20109) under accession numbers: ERS1884053 to ERS1884254, sample details including MIDS are also in Table S6. Mixed-culture sequences are available from ENA PRJEB35605 (https://www.ebi.ac.uk/ena/browser/view/PRJEB35605) with details also in Table S7. 454 data used in the time-series is available at SRA (https://trace.ncbi.nlm.nih.gov/Traces/sra/) under accession SRR097418 (11). HiSeq metagenome sequences are available at ENA project PRJEB37497 under accession numbers: ERS4415328 to ERS4415335 and the assembled metagenomes are available at IMG (https://img.jgi.doe.gov/cgi-bin/m/main.cgi) with TOIDs: 3300020141, 3300020151, 3300020159, 3300020160, 3300020161, 3300020162, 3300020172, & 3300020205 details in Table S9. Rscripts used for correlation and network analyses are available from https://github.com/rmondav/publications.

## ACKNOWLEDGEMENTS

RM, sequencing of the metagenomes, and publication supported by grants from the Malméns Stiftelsen. SG was supported by VR (grant 2012-4592), Olsson-Borgh, Knut and Alice Wallenberg Foundation (grant KAW 2013.0091), SciLifeLab Fellowship, and Kungl. Vetenskapsakademiens stiftelser (BS2017-0044). SB was supported by grants from the Swedish Research Council (VR) and the Swedish Research Council Formas. The sequencing of the Lake Erken 16S rRNA time-series dataset and chemical data was supported by the Swedish Infrastructure for Ecosystem Science (SITES) with technical assistance from Dr. Omneya Ahmed, Pilar López Hernández, Helena Enderskog, Erika Bridell and Kristiina Mustonen. Sequencing of the mixed cultures was supported by a collaborative (grant from the K&A2012-4592), Olsson-Borgh, Knut and Alice Wallenberg foundation (PI. Lars Tranvik). The funders had no role in study design, data collection and interpretation, or the decision to submit the work for publication. We gratefully acknowledge the computing resources provided by SNIC through Uppsala Multidisciplinary Centre for Advanced Computational Science (UPPMAX) under UPPNEX projects 2015047, 2016272, & 2017147 and sequencing infrastructure support from the SciLifeLab National Genomics Infrastructure.

## AUTHOR CONTRIBUTIONS

SB, SL, and RM contributed to the genomic sampling design, organization, and selection. SLG and MB designed and implemented the mixed-culture experiments. RM designed and performed the bioinformatic analyses. RM and SLG synthesized the conceptual framework which was drafted by RM in consultation with SLG, with editorial contributions from all authors.

## REFERENCES

1. Tipton L, Darcy JL, Hynson NA. 2019. A developing symbiosis: Enabling crosstalk between ecologists and microbiome scientists. Front. Microbiol. 10:1–10.

2. Koeppel A, Perry EB, Sikorski J, Krizanc D, Warner A, Ward DM, Rooney AP, Brambilla E, Connor N, Ratcliff RM, Nevo E, Cohan FM. 2008. Identifying the fundamental units of bacterial diversity: A paradigm shift to incorporate ecology into bacterial systematics. Proc. Natl. Acad. Sci. U. S. A. 105:2504–2509.

3. Rodriguez-Valera F, Martin-Cuadrado A-B, Rodriguez-Brito B, Pašić L, Thingstad TF, Rohwer F, Mira A. 2009. Explaining microbial population genomics through phage predation. Nat. Rev. Microbiol. 7:828–836.

4. Fernandez VI, Yawata Y, Stocker R. 2019. A Foraging Mandala for Aquatic Microorganisms. ISME J. 13:563–575.

5. Eriksson C, Forsberg C. 1992. Nutrient Interactions and Phytoplankton Growth during the Spring Bloom Period in Lake Erken, Sweden. Int. Rev. der gesamten Hydrobiol. und Hydrogr.

6. Bell RT, Stensdotter U, Pettersson K, Istanovics V, Pierson DC. 1998. Microbial dynamics and phosphorus turnover in Lake ErkenAdvances in Limnology 51: Lake Erken - 50 Years of Limnological Research.

7. Pettersson K, Grust K, Weyhenmeyer G, Blenckner T. 2003. Seasonality of chlorophyll and nutrients in Lake Erken - Effects of weather conditions. Hydrobiologia.

8. Haglund A-L, Törnblom E, Boström B, Tranvik L. 2002. Large differences in the fraction of active bacteria in plankton, sediments, and biofilm. Microb. Ecol. 43:232–41.

9. Lymer D, Lindström ES. 2010. Changing phosphorus concentration and subsequent prophage induction alter composition of a freshwater viral assemblage. Freshw. Biol. 55:1984–1996.

10. Burgmer T, Reiss J, Wickham S, Hillebrand H. 2010. Effects of snail grazers and light on the benthic microbial food web in periphyton communities. Aquat. Microb. Ecol. 61:163–178.

11. Eiler A, Heinrich F, Bertilsson S. 2012. Coherent dynamics and association networks among lake bacterioplankton taxa. ISME J. 6:330–42.

12. Heinrich F, Eiler A, Bertilsson S. 2013. Seasonality and environmental control of freshwater SAR11 (LD12) in a temperate lake (Lake Erken, Sweden). Aquat. Microb. Ecol. 70:33–44.

13. Eiler A, Bertilsson S. 2004. Composition of freshwater bacterial communities associated with cyanobacterial blooms in four Swedish lakes. Environ. Microbiol. 6:1228–1243.

14. Henson MW, Lanclos VC, Faircloth BC, Thrash JC. 2018. Cultivation and genomics of the first freshwater SAR11 (LD12) isolate. ISME J. 12:1846–1860.

15. Newton RJ, Jones SE, Eiler A, McMahon KD, Bertilsson S. 2011. A guide to the natural history of freshwater lake bacteria. Microbiology and molecular biology reviews: MMBR.

16. Warnecke F, Amann R, Pernthaler J. 2004. Actinobacterial 16S rRNA genes from freshwater habitats cluster in four distinct lineages. Environ. Microbiol.

17. Warnecke F, Sommaruga R, Sekar R, Hofer JS, Pernthaler J. 2005. Abundances, identity, and growth state of actinobacteria in mountain lakes of different UV transparency. Appl. Environ. Microbiol.

18. Glockner FO, Zaichikov E, Belkova N, Denissova L, Pernthaler J, Pernthaler A, Amann R. 2000. Comparative 16S rRNA analysis of lake bacterioplankton reveals globally distributed phylogenetic clusters including an abundant group of actinobacteria. Appl. Environ. Microbiol.

19. Li J, Zhang J, Liu L, Fan Y, Li L, Yang Y, Lu Z, Zhang X. 2015. Annual periodicity in planktonic bacterial and archaeal community composition of eutrophic Lake Taihu. Sci. Rep. 5:1–14.

20. Comte J, Lovejoy C, Crevecoeur S, Vincent WF. 2016. Co-occurrence patterns in aquatic bacterial communities across changing permafrost landscapes. Biogeosciences 13:175–190.

21. Garcia SL, Buck M, Hamilton JJ, Wurzbacher C, Grossart H-P, McMahon KD, Eiler A. 2018. Model Communities Hint at Promiscuous Metabolic Linkages between Ubiquitous Free-Living Freshwater Bacteria. mSphere 3:1–8.

22. Eckert EM, Baumgartner M, Huber IM, Pernthaler J. 2013. Grazing resistant freshwater bacteria profit from chitin and cell-wall-derived organic carbon. Environ. Microbiol. 15:2019–2030.

23. Neuenschwander SM, Ghai R, Pernthaler J, Salcher MM. 2017. Microdiversification in genome-streamlined ubiquitous freshwater Actinobacteria. ISME J. 12:185–198.

24. Eiler A, Mondav R, Sinclair L, Fernandez-Vidal L, Scofield DDG, Scwientek P, Martinez-Garcia M, Torrents D, McMahon KDKD, Andersson SGESGE, Stepanauskas R, Woyke T, Bertilsson S, Schwientek P, Martinez-Garcia M, Torrents D, McMahon KDKD, Andersson SGESGE, Stepanauskas R, Woyke T, Bertilsson S. 2015. Tuning fresh: radiation through rewiring of central metabolism in streamlined bacteria. ISME J. 10:1–13.

25. Kim S, Kang I, Seo J-HH, Cho J-CC. 2019. Culturing the ubiquitous freshwater actinobacterial acl lineage by supplying a biochemical ‘helper’ catalase. ISME J. 112:343640.

26. Rodriguez-R L, Tsementzi D, Luo C, Konstantinidis K. 2019. Iterative Subtractive Binning of Freshwater Chronoseries Metagenomes Identifies of over Four Hundred Novel Species and their Ecologic Preferences. bioRxiv 826941.

27. Giovannoni SJ, Cameron Thrash J, Temperton B. 2014. Implications of streamlining theory for microbial ecology. ISME J. Nature Publishing Group.

28. Tripp HJ. 2013. The unique metabolism of SAR11 aquatic bacteria. J. Microbiol. Seoul Korea 51:147–53.

29. Pernthaler J, Posch T, Šimek K, Vrba J, Pernthaler A, Glöckner FO, Nübel U, Psenner R, Amann R. 2001. Predator-Specific Enrichment of Actinobacteria from a Cosmopolitan Freshwater Clade in Mixed Continuous Culture. Appl. Environ. Microbiol.

30. Salcher MM. 2014. Same same but different: Ecological niche partitioning of planktonic freshwater prokaryotes. J. Limnol. 73:74–87.

31. Dadon-Pilosof A, Conley KR, Jacobi Y, Haber M, Lombard F, Sutherland KR, Steindler L, Tikochinski Y, Richter M, Oliver Glamp F, Suzuki MT, West NJ, Genin A, Yahel G. 2017. Surface properties of SAR11 bacteria facilitate grazing avoidance. Nat. Microbiol. 1.

32. Våge S, Storesund JE, Thingstad TF. 2013. SAR11 viruses and defensive host strains. Nature 499:E3–4.

33. Morris JJ, Lenski RE, Zinser ER. 2012. The Black Queen Hypothesis: Evolution of Dependencies through Adaptive Gene Loss. MBio.

34. Pacheco AR, Moel M, Segrè D. 2019. Costless metabolic secretions as drivers of interspecies interactions in microbial ecosystems. Nat. Commun. 10:103.

35. Pande S, Kost C. 2017. Bacterial unculurability and the formation of intercellular metabolic networks. Trends Microbiol. xx:1–13.

36. Hughes BB, Beas-Luna R, Barner AK, Brewitt K, Brumbaugh DR, Cerny-Chipman EB, Close SL, Coblentz KE, de Nesnera KL, Drobnitch ST, Figurski JD, Focht B, Friedman M, Freiwald J, Heady KK, Heady WN, Hettinger A, Johnson A, Karr KA, Mahoney B, Moritsch MM, Osterback A-MK, Reimer J, Robinson J, Rohrer T, Rose JM, Sabal M, Segui LM, Shen C, Sullivan J, Zuercher R, Raimondi PT, Menge BA, Grorud-Colvert K, Novak M, Carr MH. 2017. Long-Term Studies Contribute Disproportionately to Ecology and Policy. Bioscience 67:271–281.

37. Brown JH, Whitham TG, Morgan Ernest SK, Gehring CA. 2001. Complex species interactions and the dynamics of ecological systems: Long-term experiments. Science (80-.). American Association for the Advancement of Science.

38. Ducklow HW, Doney SC, Steinberg DK. 2009. Contributions of Long-Term Research and Time-Series Observations to Marine Ecology and Biogeochemistry. Ann. Rev. Mar. Sci. 1:279–302.

39. Franklin JF. 1989. Importance and Justification of Long-Term Studies in Ecology, p. 3–19. In Long-Term Studies in Ecology. Springer New York.

40. Philippot L, Andersson SGE, Battin TJ, Prosser JI, Schimel JP, Whitman WB, Hallin S. 2010. The ecological coherence of high bacterial taxonomic ranks. Nat. Rev. Microbiol. 8:523–9.

41. Garcia SL, Salka I, Grossart HP, Warnecke F. 2013. Depth-discrete profiles of bacterial communities reveal pronounced spatio-temporal dynamics related to lake stratification. Environ. Microbiol. Rep. 5:549–555.

42. Garcia SL, Stevens SLR, Crary B, Martinez-Garcia M, Stepanauskas R, Woyke T, Tringe SG, Andersson SGE, Bertilsson S, Malmstrom RR, McMahon KD. 2017. Contrasting patterns of genome-level diversity across distinct co-occurring bacterial populations. ISME J.

43. D ‘Autréaux B, Toledano MB. 2007. ROS as signalling molecules: Mechanisms that generate specificity in ROS homeostasis. Nat. Rev. Mol. Cell Biol. Nature Publishing Group.

44. Koonin E V, Wolf YI. 2008. Genomics of bacteria and archaea: the emerging dynamic view of the prokaryotic world. Nucleic Acids Res.2008/10/25. 36:6688–6719.

45. Bienert GP, Chaumont F. 2014. Aquaporin-facilitated transmembrane diffusion of hydrogen peroxide. Biochim. Biophys. Acta - Gen. Subj. 1840:1596–1604.

46. Zaremba-Niedzwiedzka K, Viklund J, Zhao W, Ast J, Sczyrba A, Woyke T, McMahon K, Bertilsson S, Stepanauskas R, Andersson SGE. 2013. Single-cell genomics reveal low recombination frequencies in freshwater bacteria of the SAR11 clade. Genome Biol. 14:R130.

47. Tsementzi D, Rodriguez-R LM, Ruiz-Perez CA, Meziti A, Hatt JK, Konstantinidis KT. 2019. Ecogenomic characterization of widespread, closely-related SAR11 clades of the freshwater genus “Candidatus Fonsibacter” and proposal of Ca. Fonsibacter lacus sp. nov. Syst. Appl. Microbiol. 42:495–505.

48. Moore ER, Davie-Martin CL, Giovannoni SJ, Halsey KH. 2019. Pelagibacter metabolism of diatom-derived volatile organic compounds imposes an energetic tax on photosynthetic carbon fixation. Environ. Microbiol. 00:4–6.

49. Herlemann DP, Labrenz M, Jürgens K, Bertilsson S, Waniek JJ, Andersson AF. 2011. Transitions in bacterial communities along the 2000 km salinity gradient of the Baltic Sea. ISME J. 5:1571–9.

50. Caporaso JG, Kuczynski J, Stombaugh J, Bittinger K, Bushman FD, Costello EK, Fierer N, Peña AG, Goodrich JK, Gordon JI, Huttley GA, Kelley ST, Knights D, Koenig JE, Ley RE, Lozupone CA, McDonald D, Muegge BD, Pirrung M, Reeder J, Sevinsky JR, Turnbaugh PJ, Walters WA, Widmann J, Yatsunenko T, Zaneveld J, Knight R. 2010. QIIME allows analysis of high-throughput community sequencing data. Nat. Methods 7:335–336.

51. Edgar RC. 2010. Search and clustering orders of magnitude faster than BLAST. Bioinformatics 26:2460–2461.

52. McDonald D, Price MN, Goodrich J, Nawrocki EP, DeSantis TZ, Probst A, Andersen GL, Knight R, Hugenholtz P. 2012. An improved Greengenes taxonomy with explicit ranks for ecological and evolutionary analyses of bacteria and archaea. ISME J. 6:610–8.

53. Yilmaz P, Parfrey LW, Yarza P, Gerken J, Pruesse E, Quast C, Schweer T, Peplies J, Ludwig W, Glöckner FO. 2014. The SILVA and “all-species Living Tree Project (LTP)” taxonomic frameworks. Nucleic Acids Res. 42:643–648.

54. R-Core-team. 2011. R: A Language and Environment for Statistical Computing. 2.14.1. R Foundation for Statistical Computing, Vienna, Austria.

55. RStudio. 2012. RStudio: Integrated development environment for R. 0.97.248. Boston, MA.

56. Kampstra P. 2008. Beanplot: A Boxplot Alternative for Visual Comparison of Distributions. J. Stat. Softw. 28:1–9.

57. Giraudoux P. 2012. pgirmess: Data analysis in ecology. R package version 1.5.6.

58. Weiss S, Van Treuren W, Lozupone C, Faust K, Friedman J, Deng Y, Xia LC, Xu ZZ, Ursell L, Alm EJ, Birmingham A, Cram JA, Fuhrman JA, Raes J, Sun F, Zhou J, Knight R. 2016. Correlation detection strategies in microbial data sets vary widely in sensitivity and precision. ISME J. 10:1669–1681.

59. Oksanen J, Blanchet FG, Kindt R, Legendre P, Minchin PR, O’Hara B, Simpson GL, Solymos P, Stevens MHH, Wagner H. 2013. vegan: Community ecology package. R package version 2.0-7.

60. Friedman J, Alm EJ. 2012. Inferring correlation networks from genomic survey data. PLoS Comput. Biol. 8:e1002687.

61. Kurtz ZD, Müller CL, Miraldi ER, Littman DR, Blaser MJ, Bonneau RA. 2015. Sparse and Compositionally Robust Inference of Microbial Ecological Networks. PLoS Comput. Biol. 11:e1004226.

62. Sørensen T. 1948. A method of establishing groups of equal amplitude in plant sociology based on similarity of species and its application to analyses of the vegetation on Danish commons. Biol. Skr. 5:1–34.

63. Dice LR. 1945. Measures of the Amount of Ecologic Association Between Species. Ecology 26:297–302.

64. 2014. SeqPrep. 1.2.

65. Bolger AM, Lohse M, Usadel B. 2014. Trimmomatic: A flexible trimmer for Illumina sequence data. Bioinformatics 30:2114–2120.

66. Li D, Liu CM, Luo R, Sadakane K, Lam TW. 2014. MEGAHIT: An ultra-fast single-node solution for large and complex metagenomics assembly via succinct de Bruijn graph. Bioinformatics 31:1674–1676.

67. Schmieder R, Edwards R. 2011. Quality control and preprocessing of metagenomic datasets. Bioinformatics 27:863–864.

68. Markowitz VM, Chen I-M a, Palaniappan K, Chu K, Szeto E, Grechkin Y, Ratner A, Jacob B, Huang J, Williams P, Huntemann M, Anderson I, Mavromatis K, Ivanova NN, Kyrpides NC. 2012. IMG: the Integrated Microbial Genomes database and comparative analysis system. Nucleic Acids Res. 40:D115–22.

69. Sayers EW, Barrett T, Benson D a, Bolton E, Bryant SH, Canese K, Chetvernin V, Church DM, DiCuccio M, Federhen S, Feolo M, Fingerman IM, Geer LY, Helmberg W, Kapustin Y, Landsman D, Lipman DJ, Lu Z, Madden TL, Madej T, Maglott DR, Marchler-Bauer A, Miller V, Mizrachi I, Ostell J, Panchenko A, Phan L, Pruitt KD, Schuler GD, Sequeira E, Sherry ST, Shumway M, Sirotkin K, Slotta D, Souvorov A, Starchenko G, Tatusova T a, Wagner L, Wang Y, Wilbur WJ, Yaschenko E, Ye J. 2011. Database resources of the National Center for Biotechnology Information. Nucleic Acids Res. 39:D38–51.

70. Berman HM, Westbrook J, Feng Z, Gilliland G, Bhat TN, Weissig H, Shindyalov IN, Bourne PE. 2000. The Protein Data BankNucleic Acids Research.

71. Kelley LA, Mezulis S, Yates CM, Wass MN, Sternberg MJE. 2015. The Phyre2 web portal for protein modeling, prediction and analysis. Nat. Protoc. 10:845–858.

72. Camacho C, Coulouris G, Avagyan V, Ma N, Papadopoulos J, Bealer K, Madden TL. 2009. BLAST+: architecture and applications. BMC Bioinformatics 10:421.

73. Katoh K, Kuma KI, Toh H, Miyata T. 2005. MAFFT version 5: Improvement in accuracy of multiple sequence alignment. Nucleic Acids Res. 33:511–518.

74. Niu B, Fu L, Sun S, Li W. 2010. Artificial and natural duplicates in pyrosequencing reads of metagenomic data. BMC Bioinformatics 11:187.

75. Suzek BE, Huang H, McGarvey P, Mazumder R, Wu CH. 2007. UniRef: comprehensive and non-redundant UniProt reference clusters. Bioinformatics 23:1282–8.

76. Magrane M, Consortium U. 2011. UniProt Knowledgebase: a hub of integrated protein data. Database (Oxford). 2011:bar009.

77. Kalyaanamoorthy S, Minh BQ, Wong TKF, Von Haeseler A, Jermiin LS. 2017. ModelFinder: Fast model selection for accurate phylogenetic estimates. Nat. Methods 14:587–589.

78. Kozlov AM, Darriba D, Flouri T, Morel B, Stamatakis A. 2019. RAxML-NG: a fast, scalable and user-friendly tool for maximum likelihood phylogenetic inference. Bioinformatics.

