## Supplementary_info for "Streamlined freshwater bacterioplankton *Nanopelagicales* (acI) and “*Ca*. Fonsibacter” (LD12) thrive in functional cohorts"

### Supplementary Figures

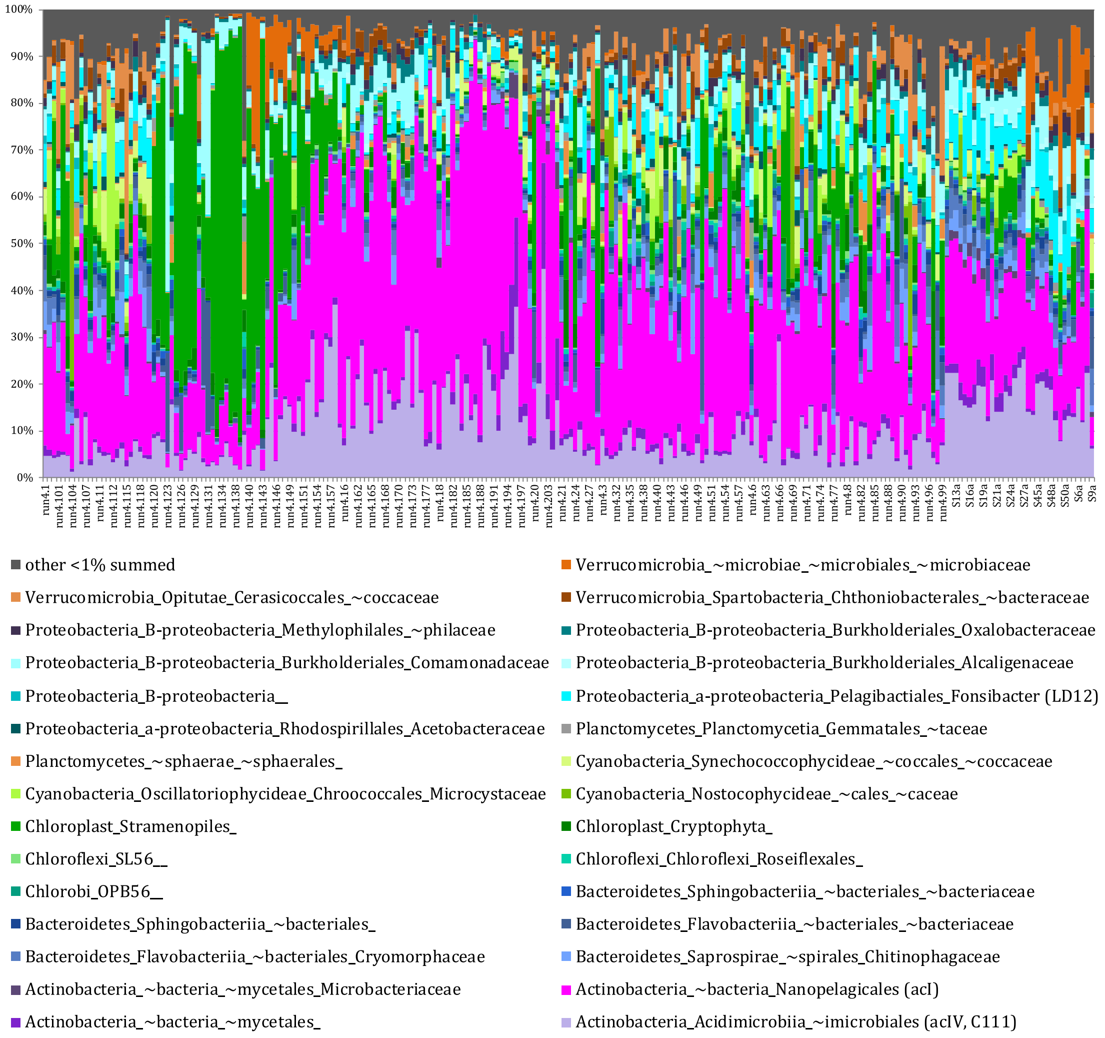

Fig S1. Relative abundance of families from community amplicon survey with total relative abundance >1.

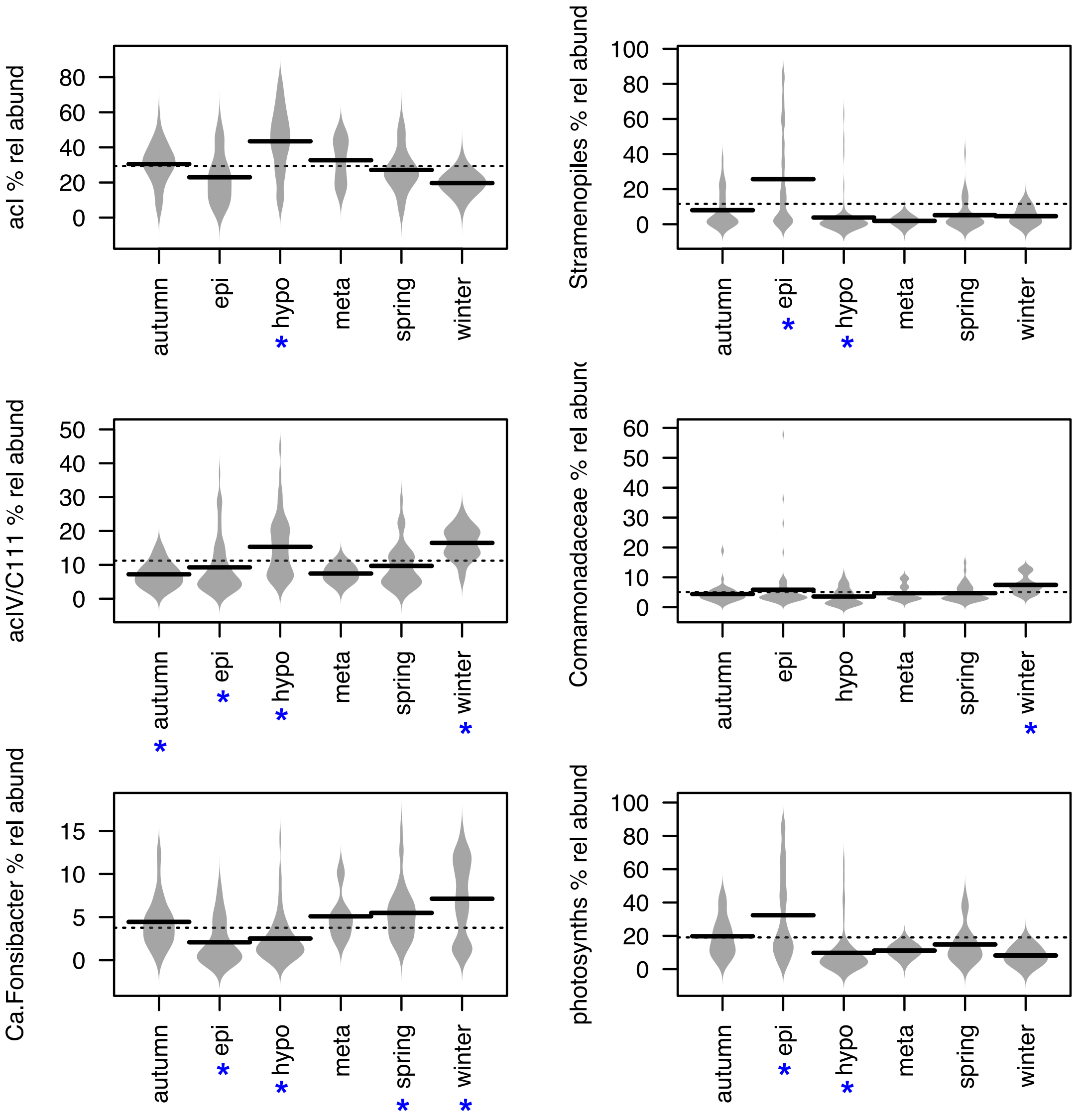

Fig S2. Associations between lake-cycle/season and dominant taxa in Lake Erken. Seasons/layers that were significantly different (KW p<0.001, KWmc p<0.001) to at least two others are denoted by asterisks.

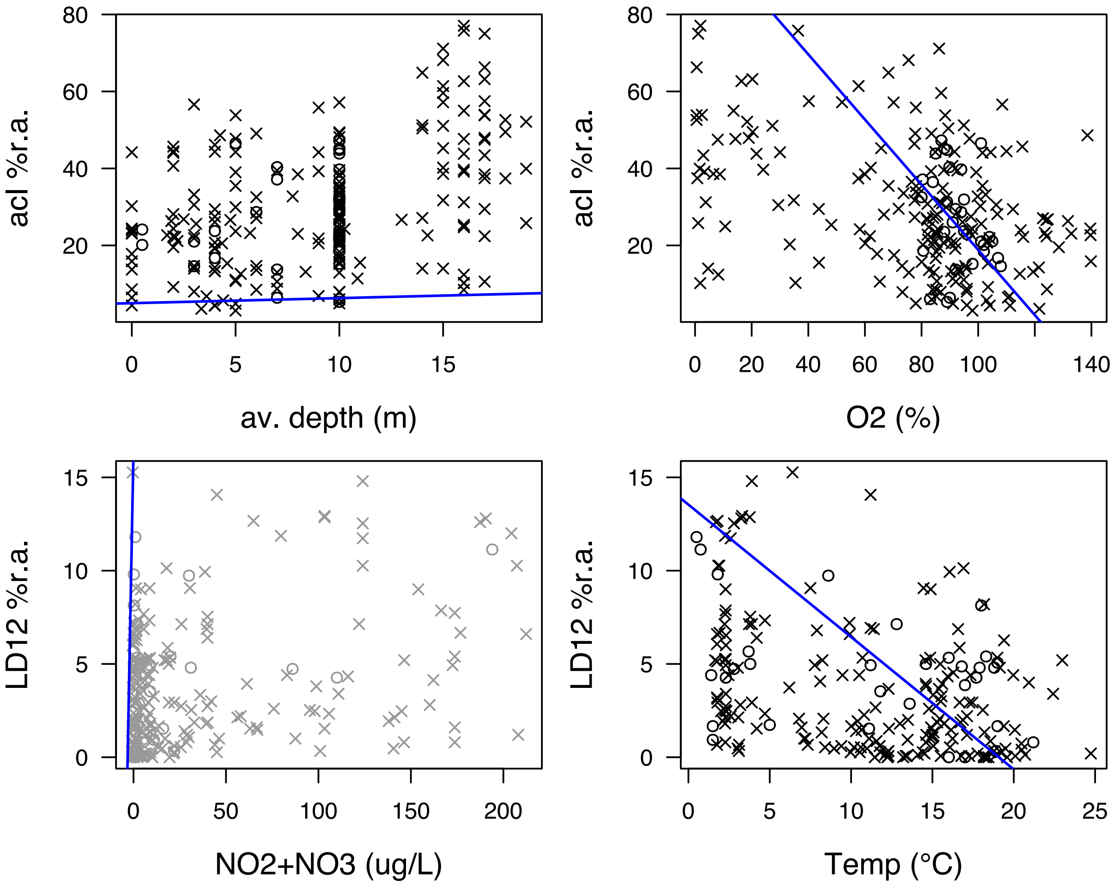

Fig S3. Correlation between taxa relative abundance and environment parameters, only showing those that have linear (Pearsons) correlation ($r >= |0.4|$, $p<0.001$). Data points with ‘x’ are 2011 to 2014 MiSeq data while ‘o’ are 454 data from 2008. The plot in grey has $r = 0.3$, $p<0.001$.

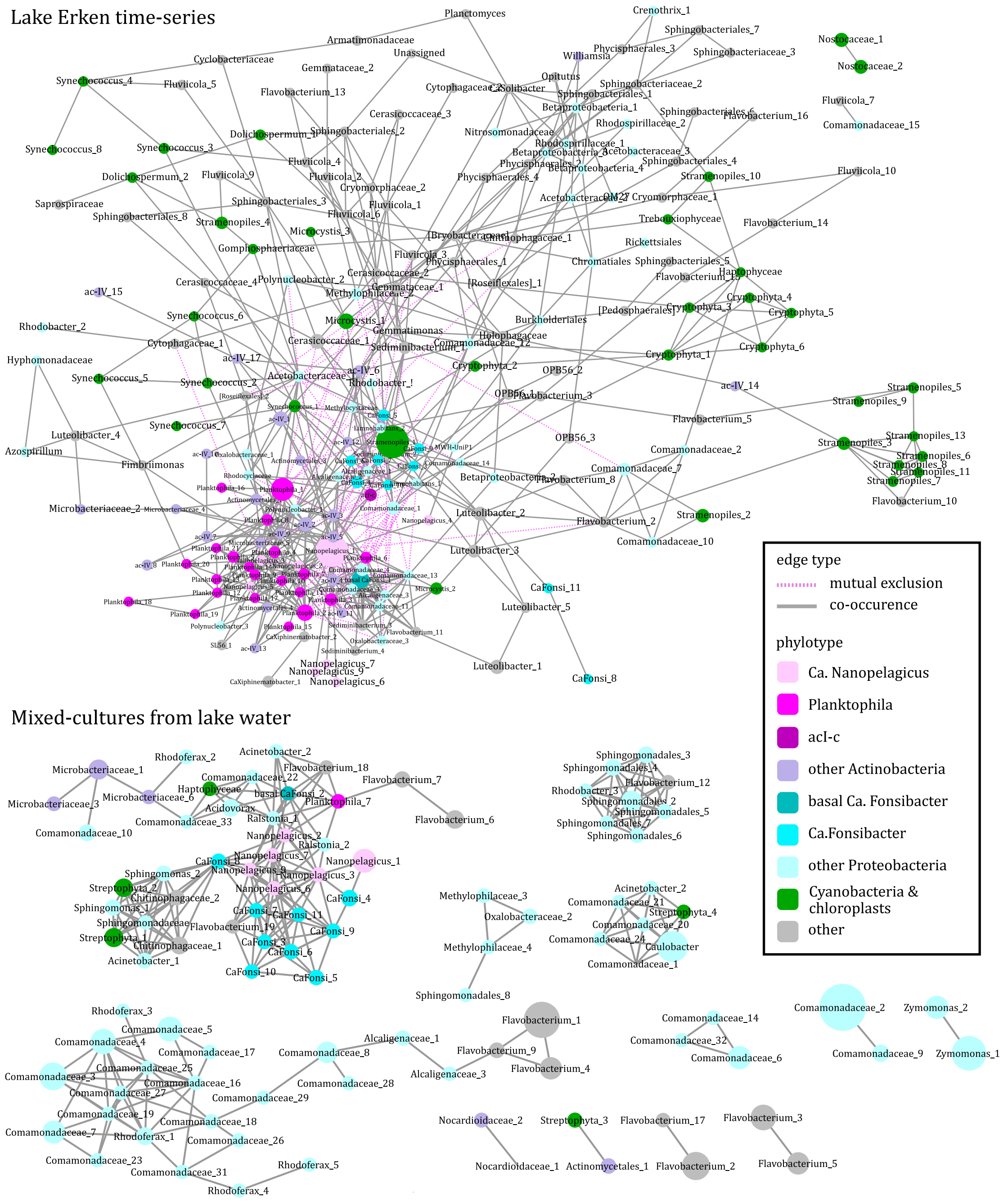

Fig S4. SSU rRNA amplicon networks based on correlations detected in at least two of three methods for the time-series and mixed-cultures

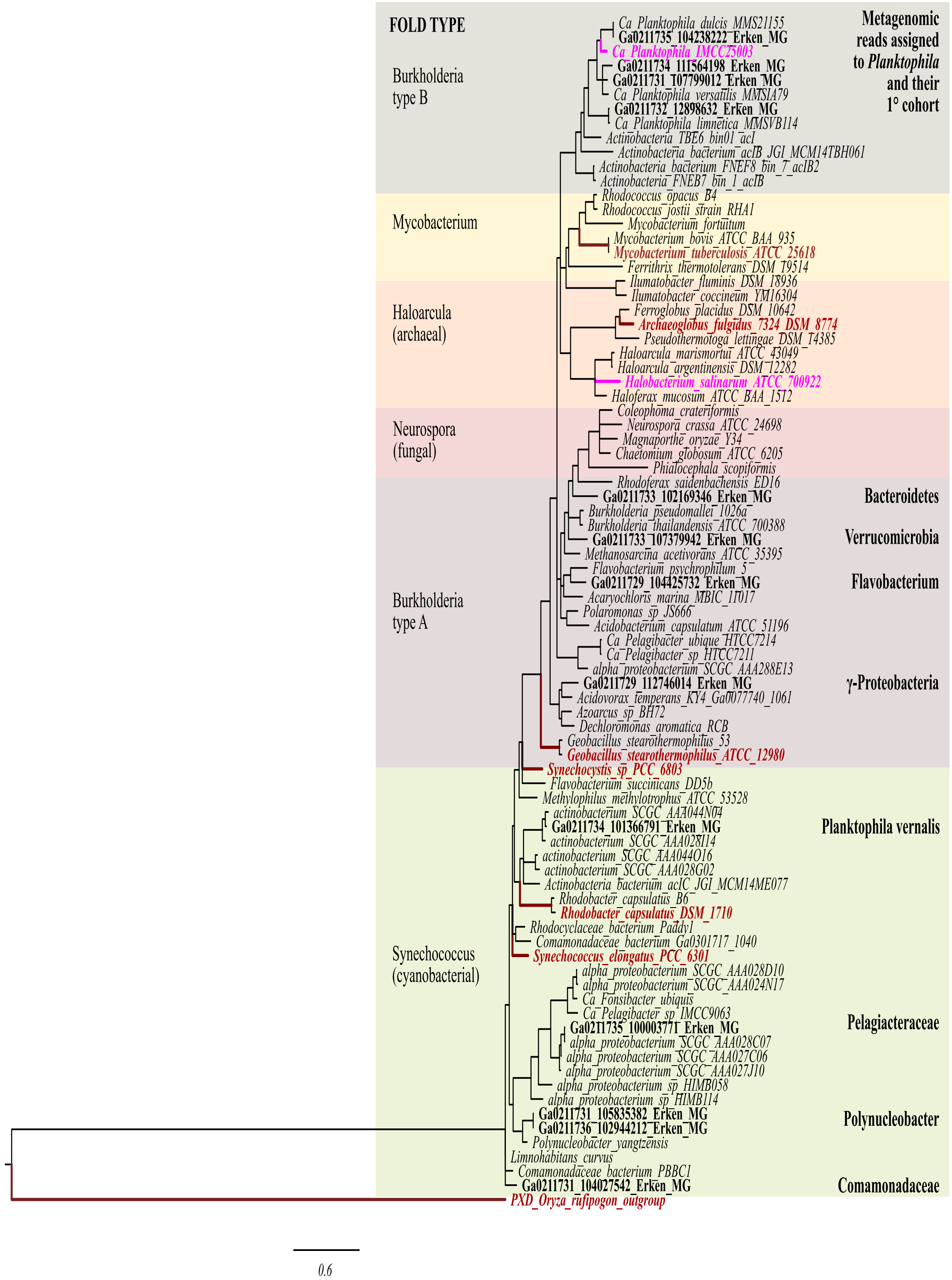

Fig. S5. RAXML phylogenetic tree of translated katG gene showing branches with over 60% support. Bold taxon names are those with assay and/or crystallography data in red for those with high function and pink for those with at least 100-fold lower activity. Metagenomic read names (starting in GA) from Lake Erken assigned to *Nanopelgicales* 1° cohort also bolded with assigned taxon listed far right.

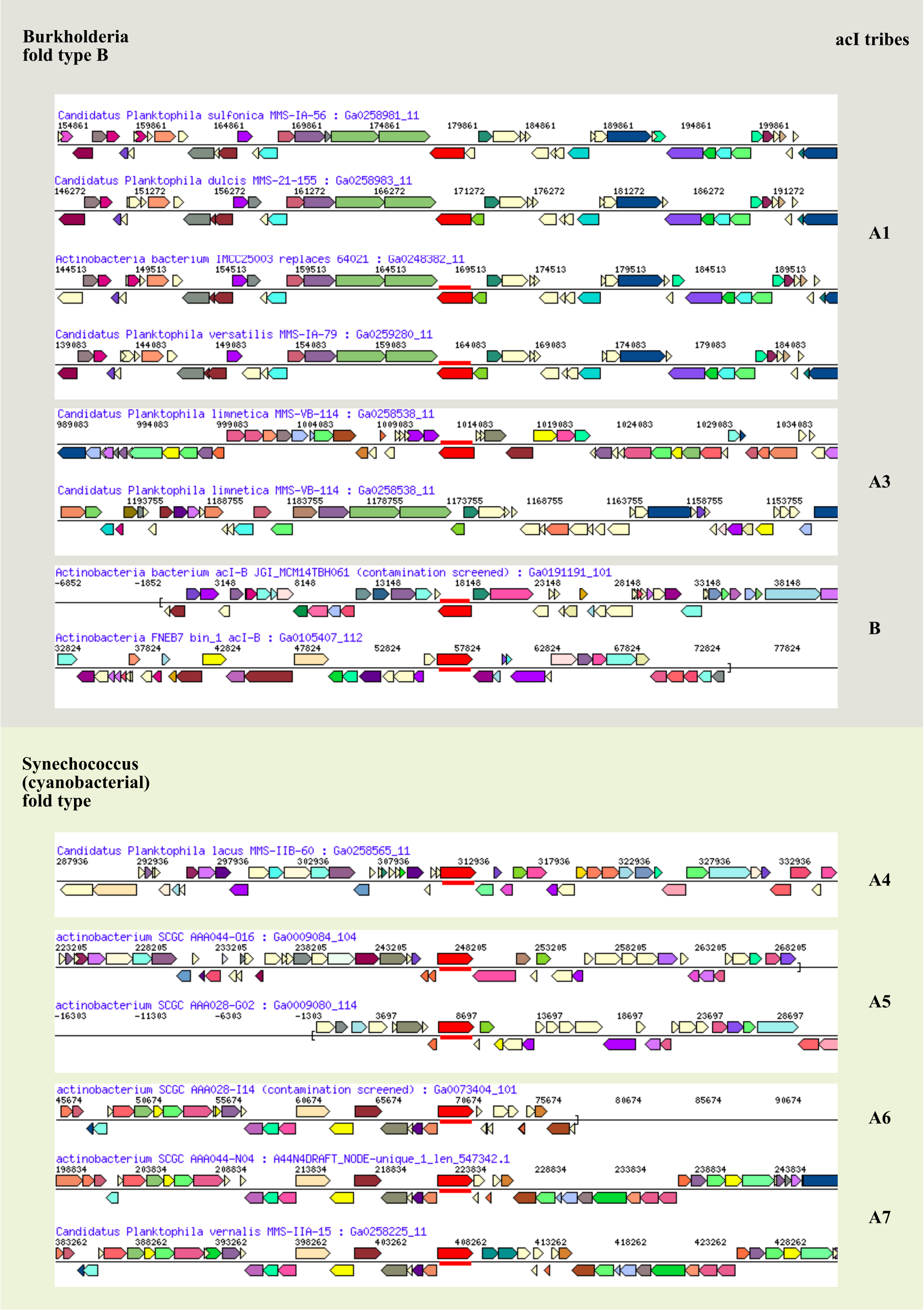

Fig. S6. Gene neighbourhoods of *katG* in *Nanopelagicales* genomes and MAGs.

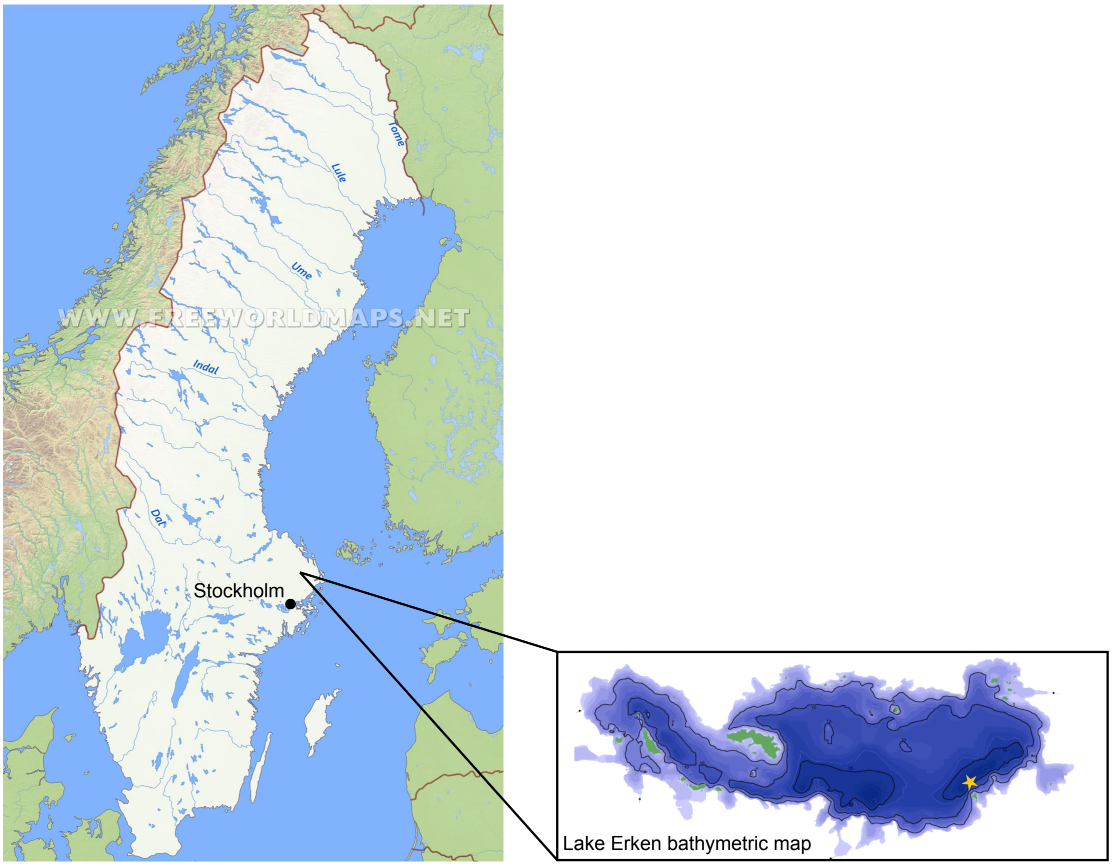

Fig S7. Location of Lake Erken in Sweden, and inset: sampling point marked with star on bathymetric map of Lake Erken.

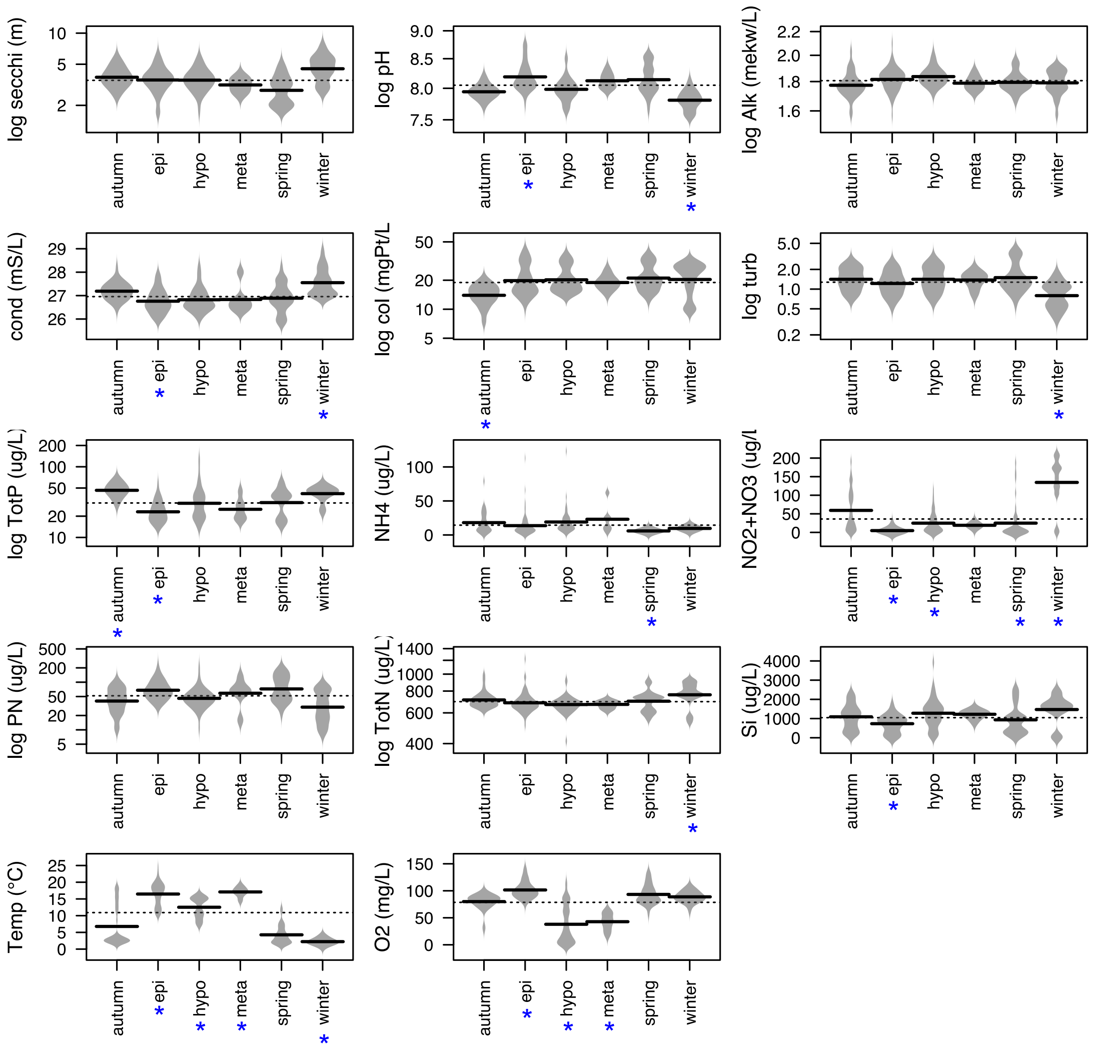

Fig S8. Associations between lake-cycle/season and environmental predictors in Lake Erken. Layers that were significantly different (KW p<0.001, KWmc p<0.001) to at least two others are denoted by asterisks below layer name.

### Supplementary Tables

Table S1. Average, maximum, minimum, and median relative abundances of families from time-series community amplicon survey

| taxonomy | Av (%) | Max (%) | Min (%) | Med (%) |
|---|---|---|---|---|
| Actinobacteria_Acidimicrobiia_~imicrobiales (acIV, C111) | 11 | 45 | 1 | 9 |
| Actinobacteria_~bacteria_~mycetales_ | 2 | 23 | 0 | 1 |
| Actinobacteria_~bacteria_Nanopelagicales (acI, ACK-M1) | 29 | 77 | 3 | 27 |
| Actinobacteria_~bacteria_~mycetales_Microbacteriaceae | 0 | 9 | 0 | 0 |
| Bacteroidetes_Saprospirae_~spirales_Chitinophagaceae | 3 | 13 | 0 | 2 |
| Bacteroidetes_Flavobacteriia_~bacteriales_Cryomorphaceae | 2 | 10 | 0 | 2 |
| Bacteroidetes_Flavobacteriia_~bacteriales_~bacteriaceae | 2 | 28 | 0 | 1 |
| Bacteroidetes_Sphingobacteriia_~bacteriales_ | 1 | 8 | 0 | 1 |
| Bacteroidetes_Sphingobacteriia_~bacteriales_~bacteriaceae | 1 | 4 | 0 | 0 |
| Chlorobi_OPB56__ | 0 | 9 | 0 | 0 |
| Chloroflexi_Chloroflexi_Roseiflexales_ | 1 | 3 | 0 | 0 |
| Chloroflexi_SL56__ | 0 | 2 | 0 | 0 |
| Chloroplast_Cryptophyta_ | 2 | 22 | 0 | 1 |
| Chloroplast_Stramenopiles_ | 12 | 88 | 0 | 2 |
| Cyanobacteria_Nostocophycideae_~cales_~caceae | 1 | 51 | 0 | 0 |
| Cyanobacteria_Oscillatoriophycideae_Chroococcales_Microcystaceae | 2 | 40 | 0 | 0 |
| Cyanobacteria_Synechococcophycideae_~coccales_~coccaceae | 2 | 17 | 0 | 1 |
| Planctomycetes_~sphaerae_~sphaerales_ | 1 | 11 | 0 | 0 |
| Planctomycetes_Planctomycetia_Gemmatales_~taceae | 1 | 8 | 0 | 0 |
| Proteobacteria_a-proteobacteria_Rhodospirillales_Acetobacteraceae | 0 | 4 | 0 | 0 |
| Proteobacteria_a-proteobacteria_Pelagibactiales_Fonsibacter (LD12) | 4 | 15 | 0 | 3 |
| Proteobacteria_B-proteobacteria__ | 1 | 5 | 0 | 0 |
| Proteobacteria_B-proteobacteria_Burkholderiales_Alcaligenaceae | 1 | 3 | 0 | 0 |
| Proteobacteria_B-proteobacteria_Burkholderiales_Comamonadaceae | 5 | 58 | 0 | 4 |
| Proteobacteria_B-proteobacteria_Burkholderiales_Oxalobacteraceae | 1 | 5 | 0 | 1 |
| Proteobacteria_B-proteobacteria_Methylophilales_~philaceae | 1 | 4 | 0 | 1 |
| Verrucomicrobia_Spartobacteria_Chthoniobacterales_~bacteraceae | 2 | 9 | 0 | 1 |
| Verrucomicrobia_Opitutae_Cerasicoccales_~coccaceae | 3 | 43 | 0 | 1 |
| Verrucomicrobia_~microbiae_~microbiales_~microbiaceae | 2 | 32 | 0 | 0 |
| other <1 summed | 8 | 31 | 1 | 7 |

Table S2. Pearson correlations (cor = Pearsons r) of taxa relative abundance and environmental parameters, p-values BH adjusted for multiple testing.

| | acI corr | acI pval | Stramen corr | Stramen pval | acIV/C111 corr | acIV/C111 pval | Comam corr | Comam pval |
|---|---|---|---|---|---|---|---|---|
| **av_depth** | 0.41 | 0.000 | -0.29 | 0.000 | 0.16 | 0.040 | -0.18 | 0.041 |
| **secchi_m** | -0.07 | 0.359 | 0.10 | 0.198 | 0.07 | 0.368 | 0.08 | 0.378 |
| **pH** | -0.09 | 0.319 | 0.10 | 0.206 | -0.08 | 0.347 | 0.05 | 0.592 |
| **Alk_mekw/L** | -0.05 | 0.549 | 0.13 | 0.122 | 0.08 | 0.368 | 0.08 | 0.378 |
| **Cond_mS/m** | -0.08 | 0.326 | -0.18 | 0.021 | 0.13 | 0.124 | 0.09 | 0.378 |
| **Col_mgPt/L** | -0.05 | 0.549 | 0.11 | 0.175 | 0.01 | 0.925 | -0.02 | 0.874 |
| **turb_FNUev26** | 0.10 | 0.274 | -0.11 | 0.175 | -0.08 | 0.368 | -0.10 | 0.339 |
| **TotP_ug/L** | 0.08 | 0.324 | -0.24 | 0.001 | 0.03 | 0.676 | 0.01 | 0.944 |
| **NH4_ug/L** | 0.11 | 0.224 | -0.06 | 0.416 | 0.14 | 0.085 | -0.07 | 0.471 |
| **NO2+NO3 ug/L** | -0.08 | 0.326 | -0.17 | 0.021 | 0.10 | 0.203 | 0.06 | 0.499 |
| **PN_ug/L** | -0.03 | 0.732 | 0.02 | 0.807 | -0.09 | 0.347 | -0.03 | 0.802 |
| **TotN_ug/L** | -0.03 | 0.693 | -0.06 | 0.405 | 0.04 | 0.594 | 0.00 | 0.995 |
| **Si_ug/L** | 0.19 | 0.013 | -0.26 | 0.001 | 0.13 | 0.124 | 0.07 | 0.471 |
| **Temp °C** | 0.09 | 0.324 | 0.21 | 0.005 | -0.03 | 0.676 | 0.01 | 0.944 |
| **O2 %** | -0.41 | 0.000 | 0.25 | 0.001 | -0.13 | 0.123 | 0.14 | 0.152 |

| | LD12 cor | LD12pval | Limnol cor | Limno pval | Polynuc cor | Polyn pval | flavo cor | flavo pval |
|---|---|---|---|---|---|---|---|---|
| **av_depth** | -0.03 | 0.709 | 0.07 | 0.437 | 0.01 | 0.962 | -0.12 | 0.240 |
| **secchi_m** | -0.03 | 0.737 | -0.12 | 0.166 | -0.02 | 0.940 | 0.03 | 0.897 |
| **pH** | -0.08 | 0.382 | 0.12 | 0.166 | 0.03 | 0.914 | 0.02 | 0.897 |
| **Alk_mekw/L** | -0.26 | 0.001 | 0.00 | 0.990 | 0.07 | 0.589 | 0.10 | 0.382 |
| **Cond_mS/m** | 0.15 | 0.070 | -0.09 | 0.359 | -0.06 | 0.616 | 0.01 | 0.970 |
| **Col_mgPt/L** | 0.04 | 0.697 | -0.07 | 0.475 | 0.02 | 0.940 | 0.00 | 0.989 |
| **turb_FNUev26** | 0.04 | 0.677 | 0.09 | 0.359 | 0.00 | 0.977 | -0.02 | 0.897 |
| **TotP_ug/L** | 0.20 | 0.008 | 0.03 | 0.689 | 0.02 | 0.940 | -0.03 | 0.897 |
| **NH4_ug/L** | -0.12 | 0.148 | 0.13 | 0.166 | 0.12 | 0.176 | -0.09 | 0.436 |
| **NO2+NO3 ug/L** | 0.33 | 0.000 | -0.23 | 0.004 | -0.11 | 0.203 | 0.02 | 0.897 |
| **PN_ug/L** | 0.02 | 0.818 | 0.06 | 0.493 | 0.01 | 0.962 | -0.02 | 0.897 |
| **TotN_ug/L** | 0.12 | 0.148 | -0.09 | 0.271 | -0.01 | 0.962 | -0.03 | 0.897 |
| **Si_ug/L** | 0.16 | 0.058 | -0.06 | 0.493 | 0.04 | 0.908 | 0.00 | 0.989 |
| **Temp °C** | -0.38 | 0.000 | 0.19 | 0.022 | 0.13 | 0.137 | -0.06 | 0.720 |
| **O2 %** | 0.06 | 0.584 | -0.04 | 0.655 | -0.07 | 0.589 | 0.09 | 0.430 |

Table S3. Statistical description of amplicon based networks

| | timeseries | mixed-culture |
|---|---|---|
| OTU count | 229 | 104 |
| $n_{eff}$ | 22 | 3 |
| sparcity | 49 | 85 |
| edges (pairs) | 625 | 252 |
| modules | 3 | 12 |
| diameter | 9 | 10 |
| av. path | 3.9 | 3.3 |
| cluster co-eff | 0.27 | 0.55 |
| density | 0.027 | 0.051 |
| av neighbours | 6 | 5 |
| phyla assort. | 0.35 | 0.21 |
| class assort. | 0.29 | 0.41 |
| order assort. | 0.29 | 0.41 |
| family assort. | 0.27 | 0.42 |
| genus assort. | 0.23 | 0.36 |
| species assort. | 0.17 | 0.34 |

Table S4. Pearson correlations (r) of OTU abundance and prevalence (% of samples) in timeseries (TS) and mixed-cultures (MC), p-values BH adjusted for multiple testing.

| All network OTUs | r | No. OTUs | p val | Test of: |
|---|---|---|---|---|
| log(sum +1) TS vs log(sum +1) MC | -0.25 | 299 | 0 | total abundance vs total abundance |
| log(av +1) TS vs log(av +1) MC | 0.58 | 299 | 0 | average abundance vs average abundance |
| log(av +1) TS vs % of MC | 0.29 | 299 | 0 | average abundance vs prevalence |
| % TS vs % of MC | -0.06 | 299 | 0.27 | prevalence vs prevalence |
| **Nanopelagicales OTUs** | **r** | **No. OTUs** | **p val** | **Test of:** |
| log(sum +1) TS vs log(sum +1) MC | 0.11 | 31 | 0.54 | total abundance vs total abundance |
| log(av +1) TS vs log(av +1) MC | 0.33 | 31 | 0.07 | average abundance vs average abundance |
| log(av +1) TS vs % of MC | 0.41 | 31 | 0.02 | average abundance vs prevalence |
| % TS vs % of MC | 0.10 | 31 | 0.61 | prevalence vs prevalence |
| **Ca. Fonsibacter OTUs** | **r** | **No. OTUs** | **p val** | **Test of:** |
| log(sum +1) TS vs log(sum +1) MC | 0.32 | 11 | 0.34 | total abundance vs total abundance |
| log(av +1) TS vs log(av +1) MC | -0.06 | 11 | 0.87 | average abundance vs average abundance |
| log(av +1) TS vs % of MC | -0.11 | 11 | 0.76 | average abundance vs prevalence |
| % TS vs % of MC | **0.82** | **11** | **0** | prevalence vs prevalence |

Table S5. Mixed cultures phylotype detection, cells colour coded as per other graphs with pinks Actinobacteria, aquas Proteobacteria, green phytoplankton, and grey all other. The number in cells is the number of phylotypes detected.

| **phylotype** | no.28 | no.88 | no.79 | no.68 | no.85 | no.57 | no.71 | no.42 | no.55 | no.69 | no.97 | no.61 | no.6 | no.38 | no.60 | no.56 | no.23 | no.82 | no.45 | no.64 | no.43 | no.34 | no.44 | no.16 | no.5 | no.26 |
|---|---|---|---|---|---|---|---|---|---|---|---|---|---|---|---|---|---|---|---|---|---|---|---|---|---|---|
| *Planktophila limnetica* | 1 | 0 | 0 | 0 | 0 | 0 | 0 | 0 | 0 | 0 | 0 | 0 | 0 | 0 | 0 | 0 | 0 | 0 | 0 | 0 | 0 | 0 | 0 | 0 | 0 | 0 |
| *Nanopelagicus* | 6 | 5 | 6 | 2 | 1 | 1 | 1 | 1 | 1 | 1 | 1 | 1 | 1 | 1 | 1 | 1 | 0 | 0 | 0 | 0 | 0 | 0 | 0 | 0 | 0 | 0 |
| *Microbacteriaceae* | 0 | 0 | 0 | 0 | 0 | 0 | 0 | 0 | 0 | 0 | 0 | 0 | 3 | 0 | 0 | 2 | 0 | 1 | 0 | 0 | 0 | 0 | 0 | 0 | 0 | 0 |
| *Actinomycetales* | 0 | 0 | 0 | 0 | 1 | 0 | 0 | 0 | 0 | 0 | 0 | 0 | 0 | 0 | 0 | 0 | 0 | 0 | 0 | 0 | 0 | 0 | 0 | 0 | 0 | 0 |
| *Nocardioidaceae* | 0 | 0 | 0 | 0 | 0 | 0 | 0 | 0 | 0 | 0 | 2 | 0 | 0 | 0 | 0 | 0 | 0 | 0 | 0 | 0 | 0 | 0 | 0 | 0 | 0 | 0 |
| *basal Ca.Fonsibacter* | 1 | 1 | 0 | 0 | 0 | 0 | 0 | 0 | 0 | 0 | 0 | 0 | 0 | 0 | 0 | 0 | 0 | 0 | 0 | 0 | 0 | 0 | 0 | 0 | 0 | 0 |
| *Ca.Fonsibacter* | 9 | 8 | 9 | 2 | 8 | 1 | 1 | 2 | 1 | 1 | 7 | 2 | 0 | 0 | 0 | 0 | 2 | 1 | 0 | 0 | 0 | 0 | 0 | 0 | 0 | 0 |
| *Acidovorax delafieldii* | 0 | 1 | 0 | 0 | 0 | 0 | 0 | 0 | 0 | 0 | 0 | 0 | 0 | 0 | 0 | 0 | 0 | 0 | 0 | 0 | 0 | 0 | 0 | 0 | 0 | 0 |
| *Acinetobacter* | 1 | 0 | 1 | 0 | 0 | 0 | 0 | 0 | 0 | 0 | 0 | 0 | 0 | 0 | 0 | 0 | 0 | 0 | 0 | 0 | 0 | 0 | 0 | 0 | 0 | 0 |
| *Alcaligenaceae* | 0 | 0 | 0 | 0 | 0 | 0 | 0 | 0 | 0 | 0 | 0 | 0 | 0 | 0 | 0 | 2 | 0 | 0 | 0 | 2 | 0 | 0 | 0 | 0 | 0 | 0 |
| *Caulobacter* | 0 | 0 | 0 | 1 | 0 | 0 | 0 | 0 | 0 | 0 | 0 | 0 | 0 | 0 | 0 | 0 | 0 | 0 | 0 | 0 | 0 | 0 | 0 | 0 | 0 | 0 |
| *Comamonadaceae* | 0 | 3 | 3 | 4 | 0 | 12 | 15 | 4 | 3 | 7 | 0 | 12 | 6 | 2 | 14 | 3 | 2 | 16 | 15 | 4 | 0 | 4 | 13 | 12 | 14 | 3 |
| *Limnohabitans* | 0 | 0 | 0 | 1 | 0 | 0 | 0 | 0 | 0 | 0 | 0 | 0 | 0 | 0 | 0 | 0 | 0 | 0 | 0 | 0 | 0 | 0 | 0 | 0 | 0 | 0 |
| *Methylophilaceae* | 0 | 0 | 0 | 0 | 0 | 0 | 0 | 0 | 3 | 0 | 0 | 0 | 0 | 0 | 0 | 2 | 0 | 0 | 0 | 0 | 0 | 0 | 1 | 1 | 1 | 0 |
| *Oxalobacteraceae* | 0 | 0 | 0 | 0 | 0 | 0 | 0 | 0 | 1 | 0 | 0 | 0 | 0 | 0 | 0 | 0 | 0 | 0 | 0 | 0 | 0 | 0 | 0 | 0 | 0 | 0 |
| *Ralstonia* | 2 | 2 | 0 | 1 | 0 | 0 | 0 | 0 | 0 | 0 | 0 | 0 | 0 | 0 | 0 | 0 | 0 | 0 | 0 | 0 | 0 | 0 | 0 | 0 | 0 | 0 |
| *Rhodoferax* | 0 | 0 | 0 | 0 | 0 | 1 | 3 | 0 | 0 | 1 | 0 | 3 | 3 | 0 | 2 | 0 | 0 | 4 | 3 | 2 | 0 | 0 | 1 | 2 | 2 | 1 |
| *Sphingomonadales* | 0 | 0 | 1 | 1 | 0 | 0 | 0 | 1 | 1 | 0 | 0 | 0 | 0 | 6 | 2 | 2 | 0 | 0 | 0 | 0 | 0 | 0 | 0 | 0 | 0 | 0 |
| *Sphingomonas* | 0 | 0 | 2 | 0 | 0 | 0 | 0 | 0 | 0 | 0 | 0 | 0 | 0 | 0 | 0 | 0 | 0 | 0 | 0 | 0 | 0 | 0 | 0 | 1 | 0 | 0 |
| *Zymomonas* | 0 | 0 | 0 | 0 | 0 | 2 | 0 | 2 | 1 | 0 | 0 | 0 | 0 | 0 | 1 | 0 | 0 | 1 | 0 | 2 | 2 | 1 | 0 | 2 | 2 | 0 |
| *Haptophyceae* | 0 | 1 | 0 | 0 | 0 | 0 | 0 | 0 | 0 | 0 | 0 | 0 | 0 | 0 | 0 | 0 | 0 | 0 | 0 | 0 | 0 | 0 | 0 | 0 | 0 | 0 |
| *Streptophyta* | 0 | 0 | 3 | 1 | 1 | 0 | 0 | 0 | 0 | 0 | 1 | 0 | 0 | 0 | 0 | 0 | 0 | 0 | 0 | 0 | 0 | 0 | 0 | 0 | 0 | 0 |
| *Chitinophagaceae* | 0 | 0 | 2 | 0 | 0 | 0 | 0 | 0 | 0 | 0 | 0 | 0 | 0 | 0 | 0 | 0 | 0 | 0 | 0 | 0 | 0 | 0 | 0 | 0 | 0 | 0 |
| *Flavobacterium* | 5 | 5 | 2 | 0 | 2 | 4 | 1 | 0 | 2 | 4 | 0 | 2 | 3 | 4 | 1 | 3 | 1 | 0 | 1 | 2 | 2 | 2 | 1 | 4 | 2 | 0 |

Table S6. Sample and accession data for MiSeq timeseries from Lake Erken. Samples in blue text denote those selected for metagenome sequencing.

| SampleID | Accession | BarcodeF | BarcodeR | sample_date | depth (m) | Stratification | Layer |
|---|---|---|---|---|---|---|---|
| **Erken.1** | ERS1884053 | TCGCCTTA | TAGATCGC | 16/07/14 | 10-20 | summer strat. | hypo |
| **Erken.10** | ERS1884054 | CAGCCTCG | TAGATCGC | 30/09/14 | 0-20 | circulation | mixed |
| **Erken.100** | ERS1884055 | TCTAGGCA | GTAAGGAG | 28/08/12 | 16-20 | summer strat. | hypo |
| **Erken.101** | ERS1884056 | TCGCCTTA | ACTGCATA | 14/02/12 | 0-20 | ice cover | mixed |
| **Erken.102** | ERS1884057 | CTAGTACG | ACTGCATA | 28/08/12 | 0-14 | summer strat. | epi |
| **Erken.103** | ERS1884058 | TTCTGCCT | ACTGCATA | 10/04/12 | 0-20 | circulation | mixed |
| **Erken.104** | ERS1884059 | GCTCAGGA | ACTGCATA | 31/07/12 | 0-8 | summer strat. | epi |
| **Erken.105** | ERS1884060 | AGGAGTCC | ACTGCATA | 29/11/11 | 0-20 | circulation | mixed |
| **Erken.106** | ERS1884061 | CATGCCTA | ACTGCATA | 23/08/11 | 0-10 | summer strat. | epi |
| **Erken.107** | ERS1884062 | GTAGAGAG | ACTGCATA | 13/11/12 | 0-20 | circulation | mixed |
| **Erken.108** | ERS1884063 | CCTCTCTG | ACTGCATA | 14/08/12 | 14-20 | summer strat. | hypo |
| **Erken.109** | ERS1884064 | AGCGTAGC | ACTGCATA | 13/09/11 | 18-20 | summer strat. | hypo |
| **Erken.11** | ERS1884075 | TGCCTCTT | TAGATCGC | 16/09/14 | 14-20 | summer strat. | hypo |
| **Erken.110** | ERS1884065 | CAGCCTCG | ACTGCATA | 9/08/11 | 0-8 | summer strat. | epi |
| **Erken.111** | ERS1884066 | TGCCTCTT | ACTGCATA | 25/09/12 | 0-20 | circulation | mixed |
| **Erken.112** | ERS1884067 | TCCTCTAC | ACTGCATA | 3/10/12 | 0-20 | circulation | mixed |
| **Erken.113** | ERS1884068 | GGTATAAG | ACTGCATA | 9/08/12 | 12-20 | summer strat. | hypo |
| **Erken.114** | ERS1884069 | CAGCTAGA | ACTGCATA | 4/07/12 | 14-20 | summer strat. | hypo |
| **Erken.115** | ERS1884070 | CCATAGCA | ACTGCATA | 31/07/12 | 12-20 | summer strat. | hypo |
| **Erken.116** | ERS1884071 | GGTATAGC | ACTGCATA | 6/09/11 | 10-14 | summer strat. | meta |
| **Erken.117** | ERS1884072 | GGTTATGC | ACTGCATA | 29/05/12 | 12-20 | summer strat. | hypo |
| **Erken.118** | ERS1884073 | TAGGCAAG | ACTGCATA | 6/09/11 | 14-20 | summer strat. | hypo |
| **Erken.119** | ERS1884074 | TTGTCCAT | ACTGCATA | 27/06/12 | 0-14 | summer strat. | epi |
| **Erken.12** | ERS1884086 | TCCTCTAC | TAGATCGC | 23/06/14 | 0-14 | summer strat. | epi |
| **Erken.120** | ERS1884076 | TCTAGGCA | ACTGCATA | 9/04/13 | 0-20 | ice cover | mixed |
| **Erken.121** | ERS1884077 | TCGCCTTA | AAGGAGTA | 15/04/13 | 0-20 | ice cover | mixed |
| **Erken.122** | ERS1884078 | CTAGTACG | AAGGAGTA | 18/04/13 | 0 | ice cover | mixed |
| **Erken.123** | ERS1884079 | TTCTGCCT | AAGGAGTA | 18/04/13 | 5.7 | ice cover | mixed |
| **Erken.124** | ERS1884080 | GCTCAGGA | AAGGAGTA | 18/04/13 | 14.25 | ice cover | mixed |
| **Erken.125** | ERS1884081 | AGGAGTCC | AAGGAGTA | 18/04/13 | 0-20 | ice cover | mixed |
| **Erken.126** | ERS1884082 | CATGCCTA | AAGGAGTA | 22/04/13 | 0 | ice cover | mixed |
| **Erken.127** | ERS1884083 | GTAGAGAG | AAGGAGTA | 22/04/13 | 4.35 | ice cover | mixed |
| **Erken.128** | ERS1884084 | CCTCTCTG | AAGGAGTA | 22/04/13 | 10.875 | ice cover | mixed |
| **Erken.129** | ERS1884085 | AGCGTAGC | AAGGAGTA | 22/04/13 | 0-8 | ice cover | mixed |
| **Erken.13** | ERS1884096 | GGTATAAG | TAGATCGC | 29/07/14 | 10-20 | summer strat. | hypo |
| **Erken.130** | ERS1884087 | CAGCCTCG | AAGGAGTA | 25/04/13 | 0 | ice cover | mixed |
| **Erken.131** | ERS1884088 | TGCCTCTT | AAGGAGTA | 25/04/13 | 3 | ice cover | mixed |
| **Erken.132** | ERS1884089 | TCCTCTAC | AAGGAGTA | 25/04/13 | 0-2 | ice cover | mixed |
| **Erken.133** | ERS1884090 | GGTATAAG | AAGGAGTA | 29/04/13 | 0-20 | circulation | mixed |
| **Erken.134** | ERS1884091 | CAGCTAGA | AAGGAGTA | 29/04/13 | 0 | circulation | mixed |
| **Erken.135** | ERS1884092 | CCATAGCA | AAGGAGTA | 29/04/13 | 3.1 | circulation | mixed |
| **Erken.136** | ERS1884093 | GGTATAGC | AAGGAGTA | 29/04/13 | 7.75 | circulation | mixed |

| | | | | | | | |
|---|---|---|---|---|---|---|---|
| **Erken.138** | ERS1884094 | TAGGCAAG | AAGGAGTA | 2/05/13 | 2.2 | circulation | mixed |
| **Erken.139** | ERS1884095 | TTGTCCAT | AAGGAGTA | 2/05/13 | 4.4 | circulation | mixed |
| **Erken.14** | ERS1884107 | CAGCTAGA | TAGATCGC | 10/02/15 | 0-20 | ice cover | mixed |
| **Erken.140** | ERS1884097 | TCTAGGCA | AAGGAGTA | 6/05/13 | 0 | circulation | mixed |
| **Erken.141** | ERS1884098 | TCGCCTTA | CTAAGCCT | 6/05/13 | 2.1 | circulation | mixed |
| **Erken.142** | ERS1884099 | CTAGTACG | CTAAGCCT | 6/05/13 | 5.25 | circulation | mixed |
| **Erken.143** | ERS1884100 | TTCTGCCT | CTAAGCCT | 2/05/13 | 0-20 | circulation | mixed |
| **Erken.144** | ERS1884101 | GCTCAGGA | CTAAGCCT | 9/05/13 | 0 | circulation | mixed |
| **Erken.145** | ERS1884102 | AGGAGTCC | CTAAGCCT | 9/05/13 | 1.9 | circulation | mixed |
| **Erken.146** | ERS1884103 | CATGCCTA | CTAAGCCT | 6/05/13 | 0-20 | circulation | mixed |
| **Erken.147** | ERS1884104 | GTAGAGAG | CTAAGCCT | 9/05/13 | 4.75 | circulation | mixed |
| **Erken.148** | ERS1884105 | CCTCTCTG | CTAAGCCT | 9/05/13 | 0-20 | circulation | mixed |
| **Erken.149** | ERS1884106 | AGCGTAGC | CTAAGCCT | 13/05/13 | 0 | summer strat. | epi |
| **Erken.15** | ERS1884118 | CCATAGCA | TAGATCGC | 24/09/14 | 18-20 | summer strat. | hypo |
| **Erken.150** | ERS1884108 | CAGCCTCG | CTAAGCCT | 13/05/13 | 4.25 | summer strat. | epi |
| **Erken.151** | ERS1884109 | TGCCTCTT | CTAAGCCT | 13/05/13 | 0-4 | summer strat. | epi |
| **Erken.152** | ERS1884110 | TCCTCTAC | CTAAGCCT | 13/05/13 | 6-20 | summer strat. | hypo |
| **Erken.153** | ERS1884111 | GGTATAAG | CTAAGCCT | 13/05/13 | 1.7 | summer strat. | epi |
| **Erken.154** | ERS1884112 | CAGCTAGA | CTAAGCCT | 16/05/13 | 0 | summer strat. | epi |
| **Erken.155** | ERS1884113 | CCATAGCA | CTAAGCCT | 16/05/13 | 2.5 | summer strat. | epi |
| **Erken.156** | ERS1884114 | GGTATAGC | CTAAGCCT | 16/05/13 | 6.25 | summer strat. | hypo |
| **Erken.157** | ERS1884115 | GGTTATGC | CTAAGCCT | 16/05/13 | 0-4 | summer strat. | epi |
| **Erken.158** | ERS1884116 | TAGGCAAG | CTAAGCCT | 16/05/13 | 6-20 | summer strat. | hypo |
| **Erken.159** | ERS1884117 | TTGTCCAT | CTAAGCCT | 20/05/13 | 0 | summer strat. | epi |
| **Erken.16** | ERS1884129 | GGTATAGC | TAGATCGC | 19/08/14 | 0-8 | summer strat. | epi |
| **Erken.160** | ERS1884119 | TCTAGGCA | CTAAGCCT | 20/05/13 | 3.35 | summer strat. | epi |
| **Erken.161** | ERS1884120 | TCGCCTTA | CTTGCTTT | 20/05/13 | 8.375 | summer strat. | epi |
| **Erken.162** | ERS1884121 | CTAGTACG | CTTGCTTT | 20/05/13 | 0-8 | summer strat. | epi |
| **Erken.163** | ERS1884122 | TTCTGCCT | CTTGCTTT | 20/05/13 | 10-20 | summer strat. | hypo |
| **Erken.164** | ERS1884123 | GCTCAGGA | CTTGCTTT | 23/05/13 | 0 | summer strat. | epi |
| **Erken.165** | ERS1884124 | AGGAGTCC | CTTGCTTT | 23/05/13 | 3.6 | summer strat. | epi |
| **Erken.166** | ERS1884125 | CATGCCTA | CTTGCTTT | 23/05/13 | 9 | summer strat. | hypo |
| **Erken.167** | ERS1884126 | GTAGAGAG | CTTGCTTT | 23/05/13 | 0-8 | summer strat. | epi |
| **Erken.168** | ERS1884127 | CCTCTCTG | CTTGCTTT | 23/05/13 | 10-20 | summer strat. | hypo |
| **Erken.169** | ERS1884128 | AGCGTAGC | CTTGCTTT | 27/05/13 | 4.6 | summer strat. | epi |
| **Erken.17** | ERS1884140 | GGTTATGC | TAGATCGC | 24/09/14 | 0-18 | summer strat. | epi |
| **Erken.170** | ERS1884130 | CAGCCTCG | CTTGCTTT | 27/05/13 | 0 | summer strat. | epi |
| **Erken.171** | ERS1884131 | TGCCTCTT | CTTGCTTT | 27/05/13 | 11.5 | summer strat. | hypo |
| **Erken.172** | ERS1884132 | TCCTCTAC | CTTGCTTT | 27/05/13 | 0-8 | summer strat. | epi |
| **Erken.173** | ERS1884133 | GGTATAAG | CTTGCTTT | 27/05/13 | 10-18 | summer strat. | hypo |
| **Erken.174** | ERS1884134 | CAGCTAGA | CTTGCTTT | 30/05/13 | 0 | summer strat. | epi |
| **Erken.175** | ERS1884135 | CCATAGCA | CTTGCTTT | 30/05/13 | 4.1 | summer strat. | epi |
| **Erken.176** | ERS1884136 | GGTATAGC | CTTGCTTT | 30/05/13 | 10.25 | summer strat. | epi |
| **Erken.177** | ERS1884137 | GGTTATGC | CTTGCTTT | 30/05/13 | 0-10 | summer strat. | epi |
| **Erken.178** | ERS1884138 | TAGGCAAG | CTTGCTTT | 30/05/13 | 12-20 | summer strat. | hypo |
| **Erken.179** | ERS1884139 | TTGTCCAT | CTTGCTTT | 3/06/13 | 0-6 | summer strat. | epi |

| | | | | | | | |
|---|---|---|---|---|---|---|---|
| **Erken.18** | ERS1884151 | TAGGCAAG | TAGATCGC | 7/07/14 | 0-20 | circulation | mixed |
| **Erken.180** | ERS1884141 | TCTAGGCA | CTTGCTTT | 8/07/13 | 0-8 | summer strat. | epi |
| **Erken.181** | ERS1884142 | TCGCCTTA | ACTTCGAC | 3/07/13 | 10-20 | summer strat. | hypo |
| **Erken.182** | ERS1884143 | CTAGTACG | ACTTCGAC | 3/07/13 | 0-6 | summer strat. | epi |
| **Erken.183** | ERS1884144 | TTCTGCCT | ACTTCGAC | 26/06/13 | 12-20 | summer strat. | hypo |
| **Erken.184** | ERS1884145 | GCTCAGGA | ACTTCGAC | 26/06/13 | 0-6 | summer strat. | epi |
| **Erken.185** | ERS1884146 | AGGAGTCC | ACTTCGAC | 18/06/13 | 10-20 | summer strat. | hypo |
| **Erken.186** | ERS1884147 | CATGCCTA | ACTTCGAC | 18/06/13 | 0-6 | summer strat. | epi |
| **Erken.187** | ERS1884148 | GTAGAGAG | ACTTCGAC | 11/06/13 | 10-20 | summer strat. | hypo |
| **Erken.188** | ERS1884149 | CCTCTCTG | ACTTCGAC | 11/06/13 | 0-8 | summer strat. | epi |
| **Erken.189** | ERS1884150 | AGCGTAGC | ACTTCGAC | 3/06/13 | 10-20 | summer strat. | hypo |
| **Erken.190** | ERS1884152 | CAGCCTCG | ACTTCGAC | 8/07/13 | 12-20 | summer strat. | hypo |
| **Erken.191** | ERS1884153 | TGCCTCTT | ACTTCGAC | 17/07/13 | 12-20 | summer strat. | hypo |
| **Erken.192** | ERS1884154 | TCCTCTAC | ACTTCGAC | 17/07/13 | 0-8 | summer strat. | epi |
| **Erken.193** | ERS1884155 | GGTATAAG | ACTTCGAC | 24/07/13 | 0-10 | summer strat. | epi |
| **Erken.194** | ERS1884156 | CAGCTAGA | ACTTCGAC | 24/07/13 | 12-20 | summer strat. | hypo |
| **Erken.195** | ERS1884157 | CCATAGCA | ACTTCGAC | 31/07/13 | 0-10 | summer strat. | epi |
| **Erken.196** | ERS1884158 | GGTATAGC | ACTTCGAC | 31/07/13 | 12-20 | summer strat. | hypo |
| **Erken.197** | ERS1884159 | GGTTATGC | ACTTCGAC | 2/08/11 | 0-10 | summer strat. | epi |
| **Erken.198** | ERS1884160 | TAGGCAAG | ACTTCGAC | 2/08/11 | 10-14 | summer strat. | meta |
| **Erken.199** | ERS1884161 | TTGTCCAT | ACTTCGAC | 2/08/11 | 14-20 | summer strat. | hypo |
| **Erken.2** | ERS1884177 | CTAGTACG | TAGATCGC | 21/07/14 | 12-20 | summer strat. | hypo |
| **Erken.20** | ERS1884167 | TCTAGGCA | TAGATCGC | 23/03/15 | 0-20 | circulation | mixed |
| **Erken.200** | ERS1884162 | TCTAGGCA | ACTTCGAC | 13/08/13 | 10-20 | summer strat. | hypo |
| **Erken.202** | ERS1884163 | CTAGTACG | TGACTTGC | 13/08/13 | 0-8 | summer strat. | epi |
| **Erken.203** | ERS1884164 | TTCTGCCT | TGACTTGC | 5/08/13 | 12-20 | summer strat. | hypo |
| **Erken.204** | ERS1884165 | GCTCAGGA | TGACTTGC | 5/08/13 | 6-12 | summer strat. | meta |
| **Erken.205** | ERS1884166 | AGGAGTCC | TGACTTGC | 5/08/13 | 0-6 | summer strat. | epi |
| **Erken.21** | ERS1884168 | TCGCCTTA | CTCTCTAT | 21/10/14 | 0-20 | circulation | mixed |
| **Erken.22** | ERS1884169 | CTAGTACG | CTCTCTAT | 4/03/15 | 0-20 | ice cover | mixed |
| **Erken.23** | ERS1884170 | TTCTGCCT | CTCTCTAT | 16/07/14 | 0-6 | summer strat. | epi |
| **Erken.24** | ERS1884171 | GCTCAGGA | CTCTCTAT | 17/03/15 | 0-20 | circulation | mixed |
| **Erken.25** | ERS1884172 | AGGAGTCC | CTCTCTAT | 9/09/14 | 0-12 | summer strat. | epi |
| **Erken.26** | ERS1884173 | CATGCCTA | CTCTCTAT | 23/06/14 | 14-20 | summer strat. | hypo |
| **Erken.27** | ERS1884174 | GTAGAGAG | CTCTCTAT | 29/07/14 | 14-20 | summer strat. | meta |
| **Erken.28** | ERS1884175 | CCTCTCTG | CTCTCTAT | 5/08/14 | 0-4 | summer strat. | epi |
| **Erken.29** | ERS1884176 | AGCGTAGC | CTCTCTAT | 31/03/15 | 0-20 | circulation | mixed |
| **Erken.3** | ERS1884188 | TTCTGCCT | TAGATCGC | 12/11/14 | 0-20 | circulation | mixed |
| **Erken.30** | ERS1884178 | CAGCCTCG | CTCTCTAT | 29/07/14 | 0-4 | summer strat. | epi |
| **Erken.31** | ERS1884179 | TGCCTCTT | CTCTCTAT | 12/08/14 | 0-4 | summer strat. | epi |
| **Erken.32** | ERS1884180 | TCCTCTAC | CTCTCTAT | 17/06/14 | 0-12 | summer strat. | epi |
| **Erken.33** | ERS1884181 | GGTATAAG | CTCTCTAT | 14/10/14 | 0-20 | circulation | mixed |
| **Erken.34** | ERS1884182 | CAGCTAGA | CTCTCTAT | 9/09/14 | 14-20 | summer strat. | hypo |
| **Erken.35** | ERS1884183 | CCATAGCA | CTCTCTAT | 26/08/14 | 14-20 | summer strat. | hypo |
| **Erken.36** | ERS1884184 | GGTATAGC | CTCTCTAT | 26/08/14 | 0-12 | summer strat. | epi |
| **Erken.37** | ERS1884185 | GGTTATGC | CTCTCTAT | 5/08/14 | 12-20 | summer strat. | hypo |

| | | | | | | | |
|---|---|---|---|---|---|---|---|
| **Erken.38** | ERS1884186 | TAGGCAAG | CTCTCTAT | 19/08/14 | 12-20 | summer strat. | hypo |
| **Erken.39** | ERS1884187 | TTGTCCAT | CTCTCTAT | 1/07/14 | 0-18 | summer strat. | epi |
| **Erken.4** | ERS1884199 | GCTCAGGA | TAGATCGC | 2/09/14 | 14-20 | summer strat. | hypo |
| **Erken.40** | ERS1884189 | TCTAGGCA | CTCTCTAT | 16/09/14 | 0-12 | summer strat. | epi |
| **Erken.41** | ERS1884190 | TCGCCTTA | TATCCTCT | 12/08/14 | 6-12 | summer strat. | meta |
| **Erken.42** | ERS1884191 | CTAGTACG | TATCCTCT | 10/06/14 | 8-20 | summer strat. | hypo |
| **Erken.43** | ERS1884192 | TTCTGCCT | TATCCTCT | 2/09/14 | 0-12 | summer strat. | epi |
| **Erken.44** | ERS1884193 | GCTCAGGA | TATCCTCT | 14/04/15 | 0-20 | circulation | mixed |
| **Erken.45** | ERS1884194 | AGGAGTCC | TATCCTCT | 9/12/14 | 0-20 | circulation | mixed |
| **Erken.46** | ERS1884195 | CATGCCTA | TATCCTCT | 12/08/14 | 12-20 | summer strat. | hypo |
| **Erken.47** | ERS1884196 | GTAGAGAG | TATCCTCT | 17/06/14 | 16-20 | summer strat. | hypo |
| **Erken.48** | ERS1884197 | CCTCTCTG | TATCCTCT | 21/04/15 | 0-20 | circulation | mixed |
| **Erken.49** | ERS1884198 | AGCGTAGC | TATCCTCT | 21/07/14 | 0-6 | summer strat. | epi |
| **Erken.5** | ERS1884210 | AGGAGTCC | TAGATCGC | 7/04/15 | 0-20 | circulation | mixed |
| **Erken.50** | ERS1884200 | CAGCCTCG | TATCCTCT | 23/08/11 | 14-20 | summer strat. | hypo |
| **Erken.51** | ERS1884201 | TGCCTCTT | TATCCTCT | 8/01/13 | 0-20 | ice cover | mixed |
| **Erken.52** | ERS1884202 | TCCTCTAC | TATCCTCT | 30/08/11 | 12-20 | summer strat. | hypo |
| **Erken.53** | ERS1884203 | GGTATAAG | TATCCTCT | 16/08/11 | 0-10 | summer strat. | epi |
| **Erken.54** | ERS1884204 | CAGCTAGA | TATCCTCT | 30/08/11 | 0-10 | summer strat. | epi |
| **Erken.55** | ERS1884205 | CCATAGCA | TATCCTCT | 19/09/12 | 0-20 | circulation | mixed |
| **Erken.56** | ERS1884206 | GGTATAGC | TATCCTCT | 6/09/11 | 0-10 | summer strat. | epi |
| **Erken.57** | ERS1884207 | GGTTATGC | TATCCTCT | 4/10/11 | 0-20 | circulation | mixed |
| **Erken.58** | ERS1884208 | TAGGCAAG | TATCCTCT | 14/08/12 | 0-8 | summer strat. | epi |
| **Erken.59** | ERS1884209 | TTGTCCAT | TATCCTCT | 13/02/13 | 0-20 | ice cover | mixed |
| **Erken.6** | ERS1884221 | CATGCCTA | TAGATCGC | 2/12/14 | 0-20 | circulation | mixed |
| **Erken.60** | ERS1884211 | TCTAGGCA | TATCCTCT | 12/10/11 | 0-20 | circulation | mixed |
| **Erken.61** | ERS1884212 | TCGCCTTA | AGAGTAGA | 27/09/11 | 0-21 | circulation | mixed |
| **Erken.62** | ERS1884213 | CTAGTACG | AGAGTAGA | 9/08/11 | 10-20 | summer strat. | hypo |
| **Erken.63** | ERS1884214 | TTCTGCCT | AGAGTAGA | 16/08/11 | 12-20 | summer strat. | hypo |
| **Erken.64** | ERS1884215 | GCTCAGGA | AGAGTAGA | 17/10/12 | 0-20 | circulation | mixed |
| **Erken.65** | ERS1884216 | AGGAGTCC | AGAGTAGA | 13/09/11 | 0-16 | summer strat. | epi |
| **Erken.66** | ERS1884217 | CATGCCTA | AGAGTAGA | 11/03/13 | 0-20 | ice cover | mixed |
| **Erken.67** | ERS1884218 | GTAGAGAG | AGAGTAGA | 11/09/12 | 0-20 | circulation | mixed |
| **Erken.68** | ERS1884219 | CCTCTCTG | AGAGTAGA | 17/04/12 | 0-18 | circulation | mixed |
| **Erken.69** | ERS1884220 | AGCGTAGC | AGAGTAGA | 24/07/12 | 0-14 | summer strat. | epi |
| **Erken.7** | ERS1884232 | GTAGAGAG | TAGATCGC | 5/08/14 | 4-12 | summer strat. | meta |
| **Erken.70** | ERS1884222 | CAGCCTCG | AGAGTAGA | 22/05/12 | 0-20 | circulation | mixed |
| **Erken.71** | ERS1884223 | TGCCTCTT | AGAGTAGA | 30/10/12 | 0-20 | circulation | mixed |
| **Erken.72** | ERS1884224 | TCCTCTAC | AGAGTAGA | 4/06/12 | 0-20 | circulation | mixed |
| **Erken.73** | ERS1884225 | GGTATAAG | AGAGTAGA | 5/09/12 | 0-20 | circulation | mixed |
| **Erken.74** | ERS1884226 | CAGCTAGA | AGAGTAGA | 9/08/12 | 0-10 | summer strat. | epi |
| **Erken.75** | ERS1884227 | CCATAGCA | AGAGTAGA | 11/07/12 | 10-20 | summer strat. | hypo |
| **Erken.76** | ERS1884228 | GGTATAGC | AGAGTAGA | 21/08/12 | 14-20 | summer strat. | hypo |
| **Erken.77** | ERS1884229 | GGTTATGC | AGAGTAGA | 27/03/12 | 0-20 | circulation | mixed |
| **Erken.78** | ERS1884230 | TAGGCAAG | AGAGTAGA | 9/05/12 | 0-20 | circulation | mixed |
| **Erken.79** | ERS1884231 | TTGTCCAT | AGAGTAGA | 14/08/12 | 8-14 | summer strat. | meta |

| | | | | | | | |
|---|---|---|---|---|---|---|---|
| **Erken.8** | ERS1884243 | CCTCTCTG | TAGATCGC | 1/07/14 | 18-20 | summer strat. | hypo |
| **Erken.80** | ERS1884233 | TCTAGGCA | AGAGTAGA | 12/06/12 | 0-20 | circulation | mixed |
| **Erken.81** | ERS1884234 | TCGCCTTA | GTAAGGAG | 21/08/12 | 0-12 | summer strat. | epi |
| **Erken.82** | ERS1884235 | CTAGTACG | GTAAGGAG | 18/01/12 | 0-20 | ice cover | mixed |
| **Erken.83** | ERS1884236 | TTCTGCCT | GTAAGGAG | 27/11/12 | 0-20 | circulation | mixed |
| **Erken.84** | ERS1884237 | GCTCAGGA | GTAAGGAG | 4/07/12 | 0-12 | summer strat. | epi |
| **Erken.85** | ERS1884238 | AGGAGTCC | GTAAGGAG | 24/04/12 | 0-20 | circulation | mixed |
| **Erken.86** | ERS1884239 | CATGCCTA | GTAAGGAG | 27/06/12 | 14-20 | summer strat. | hypo |
| **Erken.87** | ERS1884240 | GTAGAGAG | GTAAGGAG | 25/10/11 | 0-20 | circulation | mixed |
| **Erken.88** | ERS1884241 | CCTCTCTG | GTAAGGAG | 15/11/11 | 0-20 | circulation | mixed |
| **Erken.89** | ERS1884242 | AGCGTAGC | GTAAGGAG | 2/05/12 | 0-20 | circulation | mixed |
| **Erken.9** | ERS1884254 | AGCGTAGC | TAGATCGC | 10/06/14 | 0-4 | summer strat. | epi |
| **Erken.90** | ERS1884244 | CAGCCTCG | GTAAGGAG | 15/05/12 | 0-20 | circulation | mixed |
| **Erken.91** | ERS1884245 | TGCCTCTT | GTAAGGAG | 11/07/12 | 0-8 | summer strat. | epi |
| **Erken.92** | ERS1884246 | TCCTCTAC | GTAAGGAG | 24/07/12 | 16-20 | summer strat. | hypo |
| **Erken.93** | ERS1884247 | GGTATAAG | GTAAGGAG | 13/12/11 | 0-20 | circulation | mixed |
| **Erken.94** | ERS1884248 | CAGCTAGA | GTAAGGAG | 29/05/12 | 0-10 | summer strat. | epi |
| **Erken.95** | ERS1884249 | CCATAGCA | GTAAGGAG | 20/09/11 | 0-18 | circulation | mixed |
| **Erken.96** | ERS1884250 | GGTATAGC | GTAAGGAG | 18/06/12 | 12-14 | summer strat. | hypo |
| **Erken.97** | ERS1884251 | GGTTATGC | GTAAGGAG | 13/03/12 | 0-20 | ice cover | mixed |
| **Erken.98** | ERS1884252 | TAGGCAAG | GTAAGGAG | 18/06/12 | 0-10 | summer strat. | epi |
| **Erken.99** | ERS1884253 | TTGTCCAT | GTAAGGAG | 3/04/12 | 0-20 | circulation | mixed |

Table S7. Mixed-culture sequence details

| cultureID | Accession | barcodeF | barcodeR |
|---|---|---|---|
| **erken.16** | ERS4058779 | TATCCTCT | TGCCTCTT |
| **erken.23** | ERS4058786 | TATCCTCT | TAGGCAAG |
| **erken.26** | ERS4058789 | AGAGTAGA | TCGCCTTA |
| **erken.28** | ERS4058791 | AGAGTAGA | TTCTGCCT |
| **erken.34** | ERS4058797 | AGAGTAGA | AGCGTAGC |
| **erken.38** | ERS4058801 | AGAGTAGA | GGTATAAG |
| **erken.42** | ERS4058805 | AGAGTAGA | GGTTATGC |
| **erken.43** | ERS4058806 | AGAGTAGA | TAGGCAAG |
| **erken.44** | ERS4058807 | AGAGTAGA | TTGTCCAT |
| **erken.45** | ERS4058808 | AGAGTAGA | TCTAGGCA |
| **erken.5** | ERS4058768 | CTCTCTAT | TCTAGGCA |
| **erken.55** | ERS4058818 | GTAAGGAG | CAGCCTCG |
| **erken.56** | ERS4058819 | GTAAGGAG | TGCCTCTT |
| **erken.57** | ERS4058820 | GTAAGGAG | TCCTCTAC |
| **erken.6** | ERS4058769 | TATCCTCT | TCGCCTTA |
| **erken.60** | ERS4058823 | GTAAGGAG | CCATAGCA |
| **erken.61** | ERS4058824 | GTAAGGAG | GGTATAGC |
| **erken.64** | ERS4058827 | GTAAGGAG | TTGTCCAT |
| **erken.68** | ERS4058831 | ACTGCATA | TTCTGCCT |
| **erken.69** | ERS4058832 | ACTGCATA | GCTCAGGA |
| **erken.71** | ERS4058834 | ACTGCATA | CATGCCTA |
| **erken.79** | ERS4058842 | ACTGCATA | CAGCTAGA |
| **erken.82** | ERS4058845 | ACTGCATA | GGTTATGC |
| **erken.85** | ERS4058848 | ACTGCATA | TCTAGGCA |
| **erken.88** | ERS4058851 | AAGGAGTA | TTCTGCCT |
| **erken.97** | ERS4058860 | AGGTTACG | CATGCCTA |
| **erken.negC** | ERS4058862 | AAGGAGTA | CAGCCTCG |

Table S8. List of taxons identified as probable human-contamination based on literature and presence/absence in co-cultures verses timeseries data sets

| |
|---|
| k__Bacteria; p__Actinobacteria; c__Actinobacteria; o__Actinomycetales; f__Corynebacteriaceae; g__Corynebacterium; s__ |
| k__Bacteria; p__Actinobacteria; c__Actinobacteria; o__Actinomycetales; f__Corynebacteriaceae; g__Corynebacterium; s__kroppenstedtii |
| k__Bacteria; p__Actinobacteria; c__Actinobacteria; o__Actinomycetales; f__Dermacoccaceae; g__Dermacoccus; s__ |
| k__Bacteria; p__Actinobacteria; c__Actinobacteria; o__Actinomycetales; f__Micrococcaceae; g__Rothia; s__dentocariosa |
| k__Bacteria; p__Actinobacteria; c__Actinobacteria; o__Actinomycetales; f__Micrococcaceae; g__Rothia; s__mucilaginosa |
| k__Bacteria; p__Actinobacteria; c__Actinobacteria; o__Actinomycetales; f__Propionibacteriaceae; g__Propionibacterium; s__acnes |
| k__Bacteria; p__Bacteroidetes; c__Bacteroidia; o__Bacteroidales; f__Prevotellaceae; g__Prevotella; s__melaninogenica |
| k__Bacteria; p__Bacteroidetes; c__Bacteroidia; o__Bacteroidales; f__Prevotellaceae; g__Prevotella; s__nanceiensis |
| k__Bacteria; p__Bacteroidetes; c__Flavobacteriia; o__Flavobacteriales; f__Flavobacteriaceae; g__Capnocytophaga; s__ochracea |
| k__Bacteria; p__Firmicutes; c__Bacilli; o__Bacillales; f__Staphylococcaceae; g__Staphylococcus; s__aureus |
| k__Bacteria; p__Firmicutes; c__Bacilli; o__Bacillales; f__Staphylococcaceae; g__Staphylococcus; s__epidermidis |
| k__Bacteria; p__Firmicutes; c__Bacilli; o__Gemellales; f__Gemellaceae; g__; s__ |
| k__Bacteria; p__Firmicutes; c__Bacilli; o__Lactobacillales; f__Carnobacteriaceae; g__Granulicatella; s__ |
| k__Bacteria; p__Firmicutes; c__Bacilli; o__Lactobacillales; f__Lactobacillaceae; g__Lactobacillus; s__ |
| k__Bacteria; p__Firmicutes; c__Bacilli; o__Lactobacillales; f__Streptococcaceae; g__Streptococcus; s__ |
| k__Bacteria; p__Firmicutes; c__Clostridia; o__Clostridiales; f__Veillonellaceae; g__Veillonella; s__dispar |
| k__Bacteria; p__Fusobacteria; c__Fusobacteriia; o__Fusobacteriales; f__Fusobacteriaceae; g__Fusobacterium; s__ |
| k__Bacteria; p__Fusobacteria; c__Fusobacteriia; o__Fusobacteriales; f__Leptotrichiaceae; g__Leptotrichia; s__ |
| k__Bacteria; p__Proteobacteria; c__Gammaproteobacteria; o__Pasteurellales; f__Pasteurellaceae; g__Haemophilus |

Table S9. Metagenome sample details

| Name | IMG_TOIDs | ENA_Accession | TS_amp_ID | TS_amp_ERA | Description |
|---|---|---|---|---|---|
| P4710_101 | 3300020161 | ERS4415328 | Erken.50 | ERS1884200 | 23082011hypo14-20 |
| P4710_102 | 3300020172 | ERS4415329 | Erken.57 | ERS1884207 | 04102011circ0-20 |
| P4710_103 | 3300020205 | ERS4415330 | Erken.80 | ERS1884233 | 12062012circ0-20 |
| P4710_104 | 3300020141 | ERS4415331 | Erken.75 | ERS1884227 | 11072012hypo10-20 |
| P4710_105 | 3300020160 | ERS4415332 | Erken.74 | ERS1884226 | 09082012epi0-10 |
| P4710_108 | 3300020159 | ERS4415333 | Erken.36 | ERS1884184 | 26082014epi0-12 |
| P4710_201 | 3300020162 | ERS4415334 | Erken.79 | ERS1884231 | 14082012meta8-14 |
| P4710_101 | 3300020151 | ERS4415335 | Erken.27 | ERS1884174 | 29072014meta4-10 |
